# Activation of the influenza B M2 proton channel (BM2)

**DOI:** 10.1101/2024.07.26.605324

**Authors:** Zhi Yue, Jiangbo Wu, Da Teng, Zhi Wang, Gregory A. Voth

**Author notes:** Corresponding to: Gregory A. Voth. HAKUHODO Technologies Inc., Minato, Tokyo, 107-6320, Japan (Zhi Wang).

## Abstract

Influenza B viruses have co-circulated during most seasonal flu epidemics and can cause significant human morbidity and mortality due to their rapid mutation, emerging drug resistance, and severe impact on vulnerable populations. The influenza B M2 proton channel (BM2) plays an essential role in viral replication, but the mechanisms behind its symmetric proton conductance and the involvement of a second histidine (His27) cluster remain unclear. Here we perform the membrane-enabled continuous constant-pH molecular dynamics simulations on wildtype BM2 and a key H27A mutant to explore its pH-dependent conformational switch. Simulations capture the activation as the first histidine (His19) protonates and reveal the transition at lower pH values compared to AM2 is a result of electrostatic repulsions between His19 and pre-protonated His27. Crucially, we provide an atomic-level understanding of the symmetric proton conduction by identifying pre-activating channel hydration in the C-terminal portion. This research advances our understanding of the function of BM2 function and lays the groundwork for further chemically reactive modeling of the explicit proton transport process as well as possible anti-flu drug design efforts.

## Introduction

Influenza A and B viruses pose severe global health threats to humans due to their high mutation rates, emerging drug resistance, and cross-species transmission, particularly between Influenza A viruses and avian influenza (bird flu).(1, 2) Viroporins are small ion channels that facilitate virus release from infected host cells by enhancing the passive transport of ions and small molecules through the membrane, making them ideal targets for antiviral therapeutics (3-5). The matrix 2 proteins of influenza A virus (AM2) (6) and influenza B virus (BM2) (7) are proton-selective (8-16) ion channels (12, 17). Both channels are essential for viral replication (18-20) because acidifying the virion interior by proton influx through the channel is a prerequisite for releasing the virus genetic material into the host cell (21, 22).

Both channels are homotetramers (23, 24) that adopt an N_out_/C_in_ orientation in membranes (25, 26) (Fig. 1*AB*) but share little sequence homology except for an HxxxW motif (His37 and Trp41 in AM2, His19 and Trp23 in BM2, Fig. 1*C*) in the transmembrane domain. Crystallography and NMR spectroscopy have determined the closed and open structures of both channels (Table S1). In the closed-state, AM2 and BM2 pores are occluded by Val27 and Phe5 at the N-terminal (NT) side but are wide enough for water diffusion until reaching His19 and His37, respectively (Fig. 1*AB*). The presence of water in the NT-half of a closed pore, which is assumed to be a requirement for proton conductance (27), is supported by experiment (28, 29) and simulation (13-16, 29-36). It should be noted that of these numerous simulation papers only two (13, 14) have calculated the explicit channel proton conductance and activation process, and both for AM2 but not BM2. The other simulation papers have generally inferred or assumed these behaviors from looking at water within the channel and channel conformational changes, but not by also including proton transport via an explicit excess proton (or protons) in the channel and with Grotthuss proton shuttling occurring in the underlying molecular dynamics (MD) modeling algorithm.

**Figure 1.**
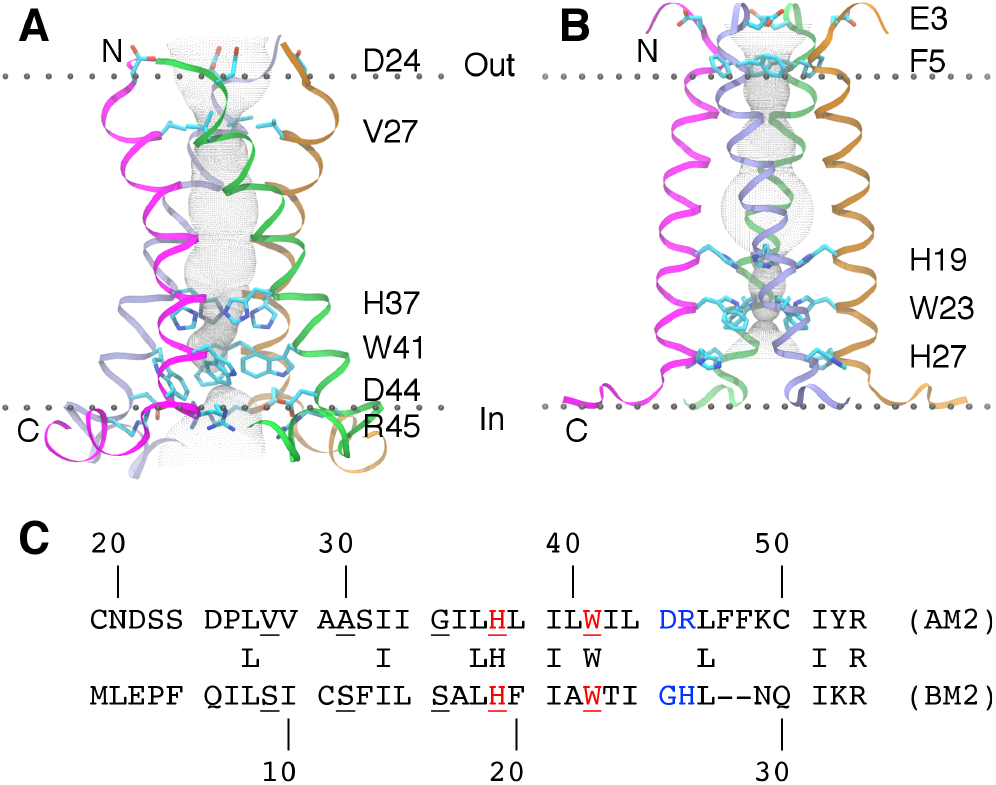
Structures and sequences of M2 proteins. Closed-state structures and crucial residues for AM2 (*A*) and BM2 (*B*). M2 tetramers (PDB IDs: 2L0J (37) and 6PVR (38) for AM2 and BM2, respectively) are shown as ribbons (ice-blue, orange, green, and magenta for chains a, b, c, and d, respectively). Residues are displayed as sticks. A gray mesh depicts the pores calculated using HOLE (39). Black dots depict the lipid head groups computed by the Positioning of Proteins in Membranes (40) (PPM) server. Membrane orientation and M2 termini are labeled. (*C*) Sequence alignment of AM2 and BM2 (UniProtKB accession numbers: P0DOF5 and B4UQM4, respectively) using the Basic Local Alignment Search Tool (41) (BLAST). Pore-lining residues are underlined. HxxxW motifs are highlighted in red. Asp44 and Arg45 in AM2, and Gly26 and His27 in BM2 are highlighted in blue.

The pores constrict at His37 in AM2 or His19 in BM2 (Fig. 1*AB*) and are respectively interrupted by densely packed Trp41 or Trp23. The pores widen after Trp41 in AM2 or Trp23 in BM2 (Fig. 1*AB*). Asp44 and Arg45 in AM2 define the C-terminal (CT) end of the pore (Fig. 1*A*), while in BM2, the pore ends correspondingly at Gly26 and a second histidine His27 (Fig. 1*B*).

In AM2, His37 is essential for the channel activity and H^+^ selectivity. Replacing His37 with Gly, Glu, Ser, or Thr removes the pH-dependent H^+^ conduction and makes the channel permeable to Na^+^ and K^+^ (42, 43). As the pH reduces in the wildtype (WT) channel, the net charge of the His37 tetrad increases from +0 to +4. AM2 activates around pH 6 (8, 10, 42, 44), where the His37 tetrad cycles between charge states of +2 and +3 based on experimental p*K*_a_s (45, 46). Upon activation, electrostatic repulsion between charged His37 residues widens the pore and opens the Trp41 gate (likely through cation–π interactions (13, 47)) for H^+^ influx (13, 14, 37, 48, 49). In addition to simulations involving explicit proton transport (13, 14), pore opening induced by charging His37 from +2 to +3 has also been captured by fixed-protonation-state (29, 30, 32, 33) (FPS) and continuous constant-pH molecular dynamics (50, 51) (CpHMD) simulations. However, it has sometime been debated if His37 is directly involved in the proton transport (PT). A “shutter” mechanism suggests that protons diffuse through a continuously hydrated open pore without changing the charge state of His37 (30, 47), while a “shuttle” mechanism argues that His37 shuttles the proton through the His37/Trp41 “bottleneck” via local conformational switch coupled with the protonation (37, 46, 48, 49, 52). Using multiscale reactive molecular dynamics (53-56) (MS-RMD) and quantum mechanics/molecular mechanics (57-59) (QM/MM) simulations, Voth and co-workers have modeled the permeation of an explicit proton through AM2 and confirmed proton “shuttling” through His37 (13-16). Trp41 also regulates the asymmetric conductance, i.e., much greater H^+^ influx when pH_out_ < pH_in_ than H^+^ efflux when pH_out_ > pH_in_ (60, 61). W41A, W41C, and W41F mutations lead to large H^+^ efflux (62). Asp44 is not strictly conserved and some naturally occurring variants have an Asn at this position (63). Mutations to Asn, Gly, Ala, Thr, Lys, and Phe make the channel more conductive (60, 64-66). Asp44 is hydrogen-bonded (HBonded) with W41 in the closed state (28, 29), and Asp44 substitutions enhance H^+^ flux by destabilizing the closed Trp41 gate (66). Asp44 also contributes to the asymmetric flux as the Cys, Ala, and Asn replacements dramatically increase H^+^ efflux when pH_out_ > pH_in_ (66). MS-RMD and QM/MM simulations have shown that the D44N mutation reduces the free energy barrier of PT between His37 and the viral interior, facilitating H^+^ efflux (67). Arg45 ion-pairs with Asp44 (28, 29) but has not been confirmed as essential for asymmetric conduction; H^+^ conduction through the R45C mutant remains asymmetric (66).

The BM2 channel displays a similar pH-dependent acidification profile (12) but differs from AM2 in several ways. First, BM2 activates and saturates at lower pH values (68). Second, BM2 has a higher activity or PT rate (68, 69). Third, the proton conductance through BM2 is symmetric, i.e., there is non-negligible H^+^ efflux when pH_out_ > pH_in_ (12, 68). Last, BM2 is insensitive to antiviral drugs amantadine (12) and rimantadine (69). In analogy to His37 in AM2, His19 is an irreplaceable pH sensor in BM2 and replacements with Cys, Ala, and Leu disrupt channel activity (12, 70-72). Trp23 is also essential to BM2 functions (70), but it has been suggested that interactions between His19 and Trp23 in BM2 differ those between His37 and Trp41 in AM2 (71). Mutants W23C and W23F are not H^+^ selective (12, 72), indicating that Trp23 is the H^+^ selectivity filter. His27 presents the most prominent structural difference in BM2. The channel activity is only partially impaired by the H27A substitution (69, 72). Replacing His27 with Arg (the AM2 equivalence) has little impact on H^+^ efflux, but mutating Gly26 to Asp (the AM2 equivalence) reduces it (68). The G26D/H27R double mutant is AM2-like and exhibits no H^+^ efflux (68), indicating that His27 is a key regulator of symmetric conductance.

Despite the electrophysiological progress, the activation mechanism of BM2 remains elusive. FPS MD simulations have suggested that the pore opens when His19 becomes triply or fully protonated (73-75). This is supported by the closed and open structures respectively solved at pH 7.5 and 4.5 (38), along with the experimental p*K*_a_s of His19 and His27 (76, 77). Gelenter et al. have combined NMR and FPS MD simulations to study the dynamics of water molecules in the channel under charge states H19^0^/H27^+1^ and H19^+4^/H27^+4^, and revealed that the higher mobility of water molecules in the open channel enables an HBonded water wire for H^+^ conduction (36). Note that the chosen charge states represent two extremes of the activation profile, so information from the transition region has not been collected. Zhang et al. have explored a broader pH range using discrete constant-pH molecular dynamics and found that activation may be triggered by the triple-protonation of the His19 tetrad (78). However, the membrane environment has not been taken into account by their employed implicit solvent generalized Born (GB) model when updating the charge states using a Monte Carlo approach. The absence of a membrane representation in the GB model, i.e., placing BM2 in a water environment, complicates the rigor of the charge-state sampling and the coupled conformational dynamics.

Here, we employed the membrane-enabled (50) hybrid-solvent (79) CpHMD technique to understand the activation of BM2 over a wide pH range. We sought to answer the following questions: (1) How does BM2 activate in response to pH reduction; and (2) how does His27 impact the channel dynamics? We confirm that BM2 opens when His19 becomes triply protonated. In addition, we observe that His27 pushes channel activation to lower pH values and increases channel hydration in the CT-portion before activation, which explains the atomistic origin of symmetric H^+^ conduction in BM2. The present work yields new insights into BM2 activation and paves the way for further investigation of the PT process (13, 14) using explicitly reactive proton transport simulations (53-56).

## Results and Discussion

### Progressive protonation of His19 and His27 tetrads

We first examined the p*K*_a_s of the protonatable residues. Microscopic p*K*_a_s assuming independent protonation were calculated for Glu3 using the Hill equation (80) (Eq. S1 of Supporting Information). For interacting residues with coupled protonation such as His19 and His27 (76, 77, 81), a four-proton model (50) (Eq. S2) was used to compute stepwise p*K*_1_/p*K*_2_/p*K*_3_/p*K*_4_. The time series of p*K*_a_ (Fig. S1) and the unprotonated fraction (S^unprot^, Figs. S2–S3) indicate convergence beyond 40 ns for all pH replicas. We thus used the 40–100 ns for analysis.

The WT simulation yields stepwise p*K*_a_s of 6.9, 6.0, 5.4, and 4.7 for His19 (Fig. 2*AD*), indicating that the tetrad stepwise stays neutral (P_0_, red line in Fig. 2*A*) at pH > 6.9, then binds the first (P_1_, orange line) and second (P_2_, brown line) protons in a pH range of 6.9–6 and 6–5.4, respectively. The triply protonated state (P_3_, green line) emerges at pH 5.4 and dominates until pH 4.7, where His19 becomes fully protonated (P_4_, blue line). The calculated p*K*_a_s are somewhat higher by < 1 pH unit than those obtained by Hong and co-workers using solid-state NMR (ssNMR) (76) (Fig. 2*D*). The dominant charge states of the His27 tetrad are P_0_, P_1_, P_2_, P_3_, and P_4_ in pH ranges of > 7.5, 7.5–6.9, 6.9–6.2, 6.2–5.3, and < 5.3, respectively (Fig. 2*BD*). Compared to the ssNMR values (77) (Fig. 2*D*), the calculated values are smaller for p*K*_1_/p*K*_4_ but higher for p*K*_2_/p*K*_3_ by *≤* 1 pH unit. The simulated sequence of protonation is H19^0^/H27^+1^ → H19^+1^/H27^+2^ → H19^+1^/H27^+3^ → H19^+2^/H27^+3^ → H19^+3^/H27^+3^ → H19^+3^/H27^+4^ → H19^+4^/H27^+4^, while the experimental sequence is H19^0^/H27^+1^ → H19^+1^/H27^+1^ → H19^+1^/H27^+3^ → H19^+1^/H27^+4^ → H19^+2^/H27^+4^ → H19^+3^/H27^+4^ → H19^+4^/H27^+4^. The major differences are 1) the simulated protonation pH range (7.5–4.7) is narrower than the experimental range (7.9–4.2), and 2) the His19 tetrad cycles between charge states P_2_ and P_3_ when His27 is triply protonated in the simulation but fully protonated in the experiment. However, the simulated p*K*_a_s of His19 are consistently smaller than those of His27, which agrees with the experiments (76, 77). The H27A mutant simulation yields stepwise p*K*_a_s of 7.0, 6.1, 5.3, and 4.6 for His19 (Fig. 2*CD*), which equal the WT values. However, the His19 protonation pH range widens to 2.4 from the WT’s 2.2, which coincides with experiments (3.6 vs. 1.9) albeit to a smaller extent.

**Figure 2.**
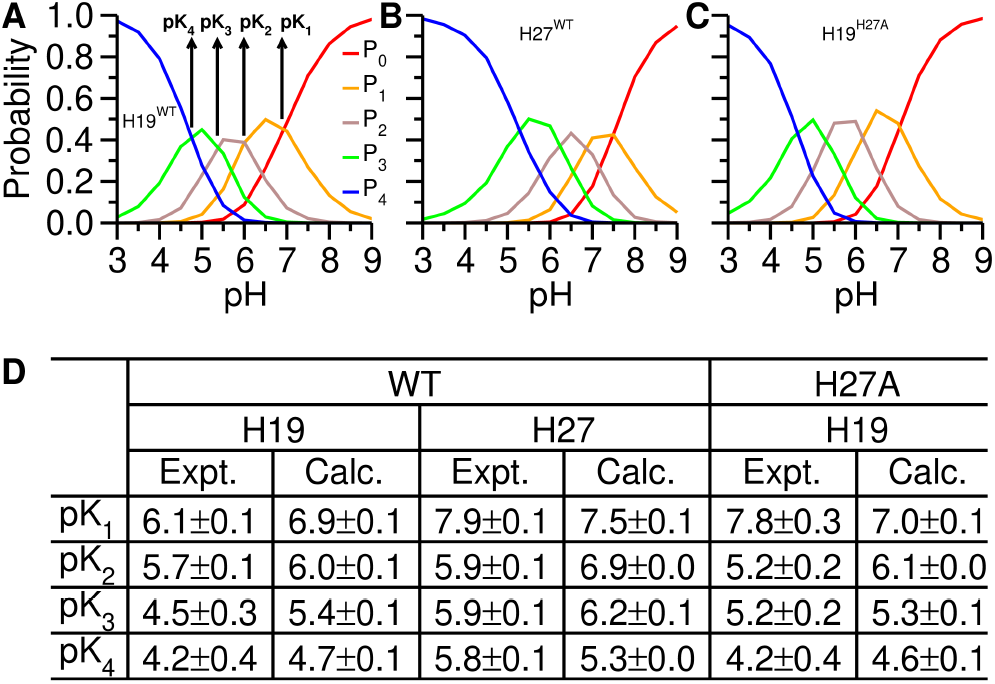
Titration of the His tetrads. Stepwise protonation of the wildtype (WT) His19 (*A*), His27 (*B*), and the H27A mutant His19 (*C*). P_0_, P_1_, P_2_, P_3_, and P_4_ represent the probabilities of binding 0 (red), 1 (orange), 2 (brown), 3 (green), and 4 (blue) protons, respectively. Stepwise p*K*_1_/p*K*_2_/p*K*_3_/p*K*_4_ are intercepts of adjacent population curves and are indicated by arrows. (*D*) Stepwise p*K*_a_s calculated using Eqs. S1–S2. The reported means and errors (standard deviations) were estimated from block analysis (block values are given in Table S6). Experimental data were taken from Hong and co-workers (76, 77, 81).

The origin of the discrepancies in p*K*_a_s is likely twofold. First, the GB model underlying the CpHMD underestimates desolvation (79), especially for deeply buried residues (82), leading to underestimated histidine p*K*_a_ shifts due to their electrostatic repulsions (79) and a narrower protonation pH range (50). Second, our simulations at 308 K used a POPC (1-palmitoyl-2-oleoyl-*sn*-glycero-3-phosphocholine) membrane while the ssNMR experiments at 243–263 K employed a virus-mimetic (VM+; POPC:POPE:sphingomyelin:cholesterol with a molar ratio of 1:1:1:1) membrane (76, 77). Under experimental conditions, BM2 is expected to be more compact and less flexible (as discussed in the subsequent section), and repulsions between histidines are reinforced, resulting in a larger p*K*_a_ differences and a wider protonation pH range.

Chen et al. studied AM2 using the CpHMD and reported stepwise p*K*_a_s of 8.3/7.1/6.2/5.7 for His37 (50), which are larger than those for His19 in BM2 by 1 pH unit or more. This raises a question: why does the histidine protonate at lower pH values in BM2? We measured the minimum distances between histidines versus pH. At pH *≥* 6 where the His19 tetrad is singly protonated or neutral, His19 residues are in close contact (a major peak at 4 Å, Fig. S4*A*). At pH *≤* 5.5 where the tetrad starts to bind more protons, the residues become separated (a second peak centers around 7 Å starts to dominate, Fig. S4*A*). This is expected given the increasing electrostatic repulsions as the charge accumulates. The separation displays a similar pH dependence in the absence of His27 (Fig. S5). The separation between His19 and His27 does not show strong pH dependence (one peak centers around 7–8 Å, Fig. S4*C*), which is not surprising based on the same-helix positioning of His19 and His27. To better assess the impact of His27, we investigated the correlation between the charge states of the His19 tetrad and that of the closest His27 residue at pH 5.5. As plotted in Fig. S4*D*, the tetrad samples more states P_1_ and P_2_ with the closest His27 residue charged but favors states P_3_ and P_4_ when the His27 is neutral. This anticorrelation (reduced repulsions with His27 favors proton binding to His19) explains the downshifted protonation region of BM2 His19 relative to that of AM2 His37.

### BM2 activates at a lower pH with a narrower transition region

We then investigated the pH-induced conformational changes of BM2. The AM2 CpHMD study revealed two well separated populations of the backbone root-mean-squared deviation (RMSD) with pH-dependent relative weights (50). However, we were unable to identify a clear pH dependence in the RMSD for BM2. The RMSD profiles display either a non-monotonic trend (WT, Fig. S6*A*) or multiple overlapping populations (H27A, Fig. S6*B*). Thus, we monitored the pore dimension along the channel axis using the quadrilateral perimeter of the pore-lining residues Phe5, Leu8, Ser12, Ser16, His19, Trp23, and His27, which is the boundary length of a polygon connecting the four C_α_ atoms. ssNMR captures changes in the perimeter of 28.3, 14.0, 9.6, 6.2, 6.2, 0.8, and – 5.6 Å for Phe5, Leu8, Ser12, Ser16, His19, Trp23, and His27, respectively, when going from the closed (PDB ID: 6PVR) to the open (PDB ID: 6PVT) state (Table S2). This indicates that BM2 undergoes a rigid scissor motion with the hinge at Trp23 upon activation in the experiment (38). While in our simulation, all perimeters except at Phe5 rapidly increase as the pH reduces from 6.5 to 4.5 (Fig. 3*A*). The closer to the CT, the larger the increase (4.0, 4.2, 5.8, 7.1, 10.0, and 12.8 Å for Leu8, Ser12, Ser16, His19, Trp23, and His27, respectively). The ssNMR structures, determined in rigid gel-phase POPE (1-palmitoyl-2-oleoyl-*sn*-glycero-3-phosphoethanolamine) membranes at 280–290 K, contrast with BM2’s dynamics in fluid-phase POPC membranes at 308 K in our simulations.

**Figure 3.**
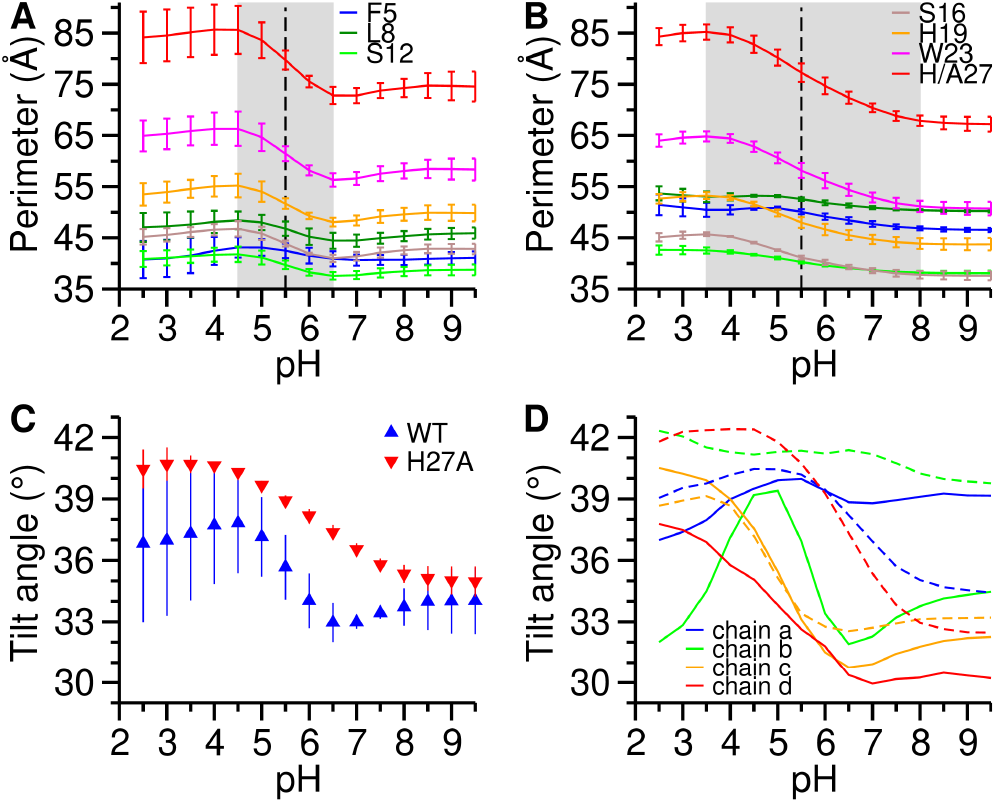
pH-dependent conformational change. Quadrilateral perimeters of the C_α_ atoms of pore-lining residues in the WT (*A*) and H27A mutant (*B*) simulations. Pore-lining residues include: Phe5 (blue), Leu8 (dark green), Ser12 (green), Ser16 (brown), His19 (orange), Trp23 (magenta), and His/Ala27 (red). Means over the simulation time were plotted versus pH values and errors were standard deviations estimated from block analysis (same below). Gray boxes and vertical black lines mark the transition regions and midpoints, respectively. Midpoints were computed by fitting relative changes to the Hill equation (80) (Eq. S1). A perimeter comparison between the WT and H27A is given in Fig. S7. The distributions are displayed in Figs. S8 (WT) and S9 (H27A). (*C*) Average tilt angle between the C-terminal helices (residues 15–28) and the N-terminal helical bundle (residues 7–14) in the WT (blue up triangle) and H27A mutant (red down triangle). Note that the calculations were insensitive to atom selections (Fig. S14). (*D*) Tilt angle between each C-terminal helix (blue, green, orange, and red for chains a, b, c, and d, respectively) and the N-terminal helical bundle in the WT (solid line) and H27A mutant (dashed line). For clarity, error bars are not shown (displayed in Fig. S15).

To assess the impact of lipid composition on channel behavior, we performed μs-long FPS simulations for the closed (H19^0^/H27^+1^) and open (H19^+3^/H27^+4^) states in POPC and VM+ bilayers at 308 K, and in POPE bilayers at 280 K. Note that VM+ membranes at 308 K are in a fluid phase (83) and POPE membranes at 280 K are in a gel phase (84-87). We did not run VM+ simulations between 243–263 K due to known issues with the TIP3P water model (88, 89) below the freezing point (90). In the FPS simulations with a gel-phase POPE bilayer, it is not surprising that the perimeters of the pore-lining residues stay close to the ssNMR values, despite a relatively constricted pore between Ser12 and Trp23 (Fig. S10). Regarding the fluid-phase simulations, the time series of RMSD indicates that BM2 is more stable in VM+ membranes than in POPC membranes in either the closed (Fig. S11, red vs. orange lines) or open (Fig. S11, blue vs. green lines) states. Consistent with the WT CpHMD simulation, all the perimeters increase upon activation in the FPS simulations with a POPC bilayer (Fig. S12, solid lines), with larger increases observed closer to the CT. By contrast, in the FPS simulations with a VM+ bilayer, the perimeters increase at His19 and above but decrease at His27, while remaining nearly unchanged at Trp23 (Fig. S12, dashed lines). This pattern corresponds with the ssNMR structures and indicates that BM2 undergoes a rigid scissor motion with the hinge at Trp23 in fluid-phase VM+ membranes. This is likely due to the immobilization of BM2 in cholesterol-rich VM+ membranes, which has also been observed for AM2 in VM+ membranes at physiological temperatures using ssNMR (83).

The WT CpHMD simulation shows that the His19 tetrad takes two protons within the transition region pH 6.5–4.5 and oscillates between charge states P_2_ and P_3_ at the transition midpoint pH 5.5 (Fig. 2*D*). This supports earlier MD simulations (73, 78) and ssNMR experiments (38, 76, 77), confirming BM2 activation upon triple-protonation of His19. Notably, the simulated activation midpoint is lower than that of AM2 (pH 6.0) (50), aligning with electrophysiological observations that BM2 activates at lower pH values (68). The H27A simulation reveals notable variances, with a broader transition region of pH 3.5–8 in the perimeter profiles (Fig. 3*B*). However, the transition midpoint remains the same (Fig. 3*B*). Otomo et al. found that the H27A mutation reduced the maximum H^+^ conductance without altering the transition midpoint (72). Despite a 1 pH unit difference between the calculated (5.5) and measured (6.4) midpoints, likely due to the systematic shift caused by the GB mode and different experimental lipid environments, our simulations concur with the experiments regarding the mutant’s impact. Like the WT, all mutant perimeters progressively increase as the pH is lowered. But there are sensible differences. Compared to the WT, H27A has a wider pore at Phe5 and Leu8 at all pH values but narrows at Ser16 and below at pH > 4 (Figs. 3*AB* and S7). This is likely due to the absence of electrostatic repulsions between His19 and His27 in the mutant. As the WT channel stays closed (Fig. 3*A*) when His27 protonates (Fig. 2*BD*) at pH > 6.5, it is expected that the CT shrinkage becomes significant at pH > 6.5 in H27A (Fig. S7). The contrasting trends of expansion at the NT and shrinkage at the CT suggest a rigid motion in BM2 helices during activation.

Another feature is the tilt angle between the CT-half helix (residues 7–14 and 25–32 for AM2 and BM2, respectively) and the NT-half helical bundle (residues 15–28 and 33–46 in AM2 and BM2), which has proven useful for probing AM2 activation (91). Like in the AM2 CpHMD simulation (50), the NT helical bundle’s tilt relative to the membrane normal is less sensitive to pH in the BM2 simulation (Fig. S13). The average tilt angle between the CT-helices and the NT-bundle slightly increases from around 34º to 37º when BM2 activates at low pH values (Fig. 3*C*, blue up triangles). ssNMR structures record an increase from 20º (PDB ID: 6PVR) to 22º (PDB ID: 6PVT). In contrast, AM2 sees an increase from 21º (PDB ID: 3LBW) to 37º (PDB ID: 3C9J) in crystallography and from 23º to 44º in the CpHMD simulation (50). Our BM2 simulation aligns with these findings, showing a smaller tilt angle increase upon activation compared to AM2. The H27A mutant displays similar behavior but reports a larger CT tilt at low pH values (40º; Fig. 3*C*, red down triangles). Overall trends of tilting agree with the perimeter ones (the same transition region and midpoint). But inspecting tilt angles of each helix show asynchronous tilting in WT (considerable increases in chains c and d as pH reduces, minor decrease in chain a, and a big decrease after an initial increase in chain b; Fig. 3*D*, solid lines), whereas in H27A, tilting is synchronous (a uniform increase across all four chains; Fig. 3*D*, dashed lines).

### H27 increases the C-terminal hydration

We now turn to channel hydration and count water molecules along the channel axis (N_Water_) in both the WT and the H27A mutant. The WT hydration remains unaltered until pH 6.5, then starts to increase at pH 6, where His19 takes the second H^+^ (Fig. 2*AD*), and continues rising as the pH further reduces (Fig. S16*A*). The change in hydration varies by location. Going from the closed state at pH 8 (His19^0^/His27^0^) to the open state at pH 4 (His19^+4^/His27^+4^), the channel experiences a greater increase in hydration in the CT-half than in the NT-half (Fig. S17*A*), consistent with the perimeter-based channel expansion (Fig. 3*A*). The increase in hydration is pronounced between His27 and Ser16 (–13 Å *≤* Z *≤* 1 Å) and maximizes (6 H_2_O molecules) around Trp23 (Z = –7 Å).

The H27A hydration profile displays a similar trend of increasing channel hydration with decreasing pH, particularly in the CT-half between Ala27 and Ser16 (Figs. S16*B* and S17*B*). However, there is a notable difference: the closed WT contains 1–2 more H_2_O molecules per 2-Å layer between His27 and Ser12 (–9 Å *≤* Z *≤* 3 Å) than H27A (Fig. 4*A*, left panel). When His19 is singly or doubly protonated, the WT channel also exhibits increased hydration in this region (Fig. 4*A*, middle and right panels). This suggests that His27 contributes to increased hydration in the CT-portion before activation. In AM2, Asp44 and Arg45 (equivalent to BM2’s Gly26 and His27) form interhelical salt-bridges in the closed state at high pH values (50), which likely seals the CT-entrance and blocks water entry until His37’s triple-protonation from the NT side leads to channel expansion, resulting in asymmetric H^+^ conduction. This is supported by MS-RMD simulations of the WT (15) and D44N mutant (67) of AM2, which found increased hydration by about 2 H_2_O molecules below Trp41 before His37 becomes triply protonated. The H_2_O network altered by the D44N mutation explains the reduced free energy barrier for protonating His37 from the interior (67) and the symmetric H^+^ conduction in the mutant (66). While in BM2, lacking such salt-bridges, His27 protonates before His19, drawing water molecules into the channel and enabling H^+^ access to His19 from the CT side, explaining the symmetric H^+^ conduction in BM2 (Fig. 4*B*). This is supported by mutagenesis studies: the H27R mutant mimics WT, the G26D mutant reduces the H^+^ efflux, and the AM2-like G26D/H27R double mutant blocks it (68). The H^+^ efflux is not blocked in the G26D mutant likely because His27 side-chains are not bulky enough to effectively block water entry if ion-paired with Asp26. These findings suggest that the H^+^ efflux in M2 proteins requires a precise geometrical side-chain arrangement at the CT-entrance. This result may also present an opportunity for a drug target.

**Figure 4.**
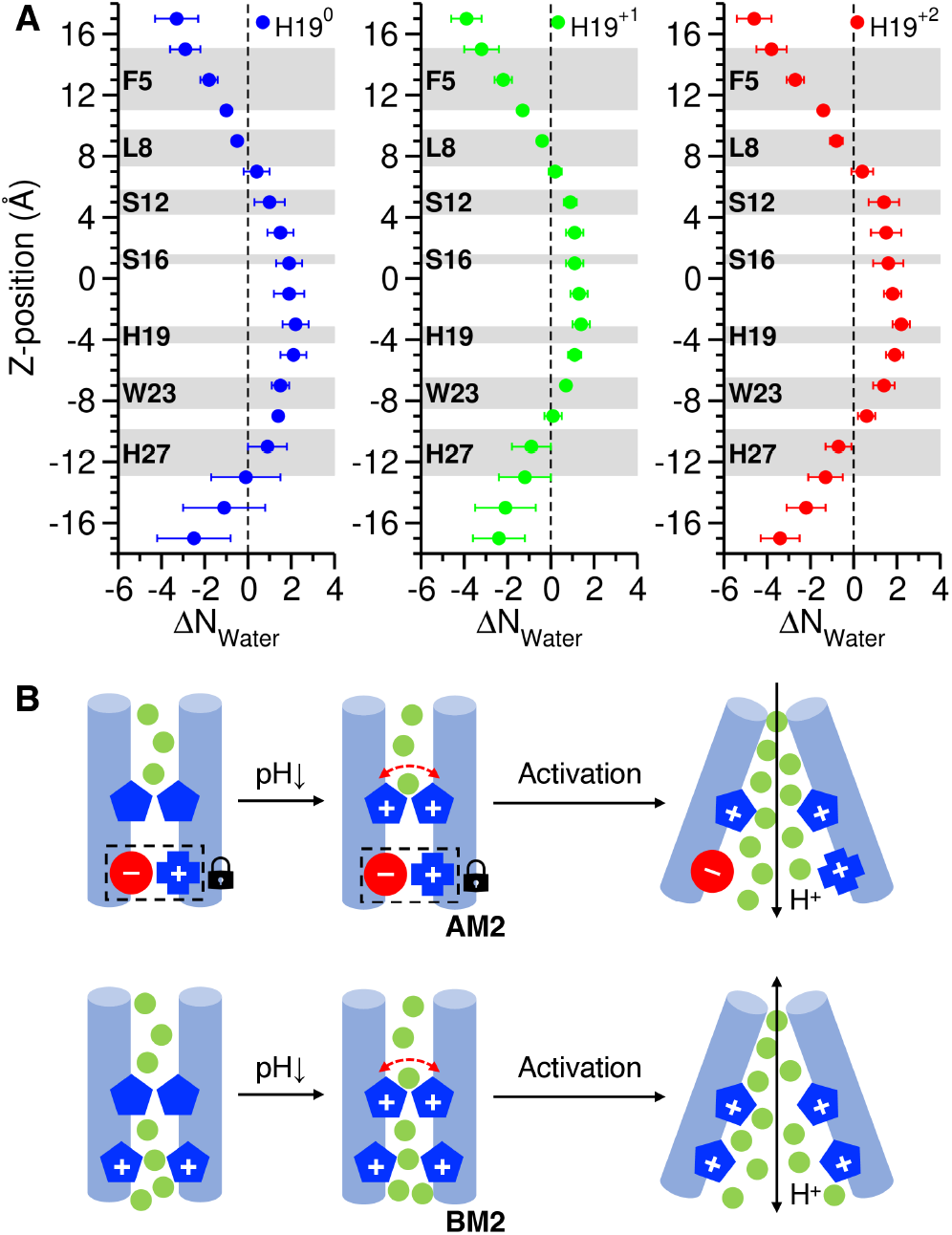
pH-dependent channel hydration. (*A*) Differences in channel hydration between the WT and H27A mutant (ΔN_Water_) when His19 is neutral (left), singly protonated (middle), and doubly protonated (right). Errors were calculated through error propagation. Gray boxes indicate the side-chain center-of-mass (COM) of the pore-lining residues (centroid and halved box width respectively represent the mean and standard deviation). Vertical dashed lines mark ΔN_Water_ of 0. The COMs were computed using heavy atoms and projected to the channel axis, with the four Ala17 C_α_ atoms centered at the origin. (*B*) Model depicting proton transport (PT) directionality in AM2 (top) and BM2 (bottom). Blue pentagons, red circles, blue crosses, and green circles represent the M2 histidines, AM2 Asp44, AM2 Arg45, and H_2_O, respectively. Red dashed arches denote electrostatic repulsions between protonated AM2 His37 or BM2 His19. In the closed-stat AM2, only the NT-half of the channel is hydrated as Asp44–Arg45 salt-bridges block H_2_O accessibility from the CT-entrance, making the PT asymmetric. While in the closed-state BM2, protonated His27 opens the CT-entrance and provides H_2_O accessibility, leading to symmetric PT.

## Conclusions

In summary, we have employed membrane-enabled constant-pH MD simulations to investigate the conformational transition of BM2 over a broad pH range. Aligning with prior experiments, BM2 activates with the triple-protonation of His19, but activation is shifted to lower pH values compared to influenza AM2, which can be attributed to electrostatic repulsion between His19 and the early protonated His27. Importantly, we identified the essential role of His27 in the pre-activation hydration of the CT-portion of the channel, which provides an atomic-level explanation for BM2’s symmetric H^+^ conduction. In addition, we observed that the lipid composition has a significant impact on the BM2 conformational dynamics, favoring the rigid scissor motion upon activation in membranes with reduced fluidity. This work therefore yields new insight into BM2 activation and paves the way for in-depth explicit proton transport modeling of the PT process, as well as possible target identification for drug design.

### Experimental procedures

We performed membrane-enabled (50) hybrid-solvent (79) CpHMD simulations on a closed ssNMR structure (PDB ID: 6PVR (38)) using the CHARMM program (92). Consistent with our influenza AM2 studies (13-16, 67), a POPC membrane was used. All simulations used the CHARMM22/CMAP (93-95) force field, the CHARMM36 model (96-98), the CHARMM-modified (89) TIP3P model (88), and the Beglov and Roux’s Lennard-Jones model (99) with NBFIX terms (100, 101) to represent the proteins, lipids, waters, and ions, respectively. Systems were stepwise equilibrated following an established protocol (102, 103). In the production run, the pH-based replica exchange protocol (79) was employed to speed up the sampling of BM2 conformations and the protonation states of Glu3, His19, and His27. A total of 15 replicas, ranging from pH 2.5 to 9.5, were run under 308.15 K and 1 atmosphere pressure for 100 ns each, resulting in an aggregate sampling time of 1.5 μs. The simulation of the H27A mutant was also initiated from the closed structure. Details of simulation settings and analysis protocols can be found in the Supporting Information.

## Data availability

All data required for the conclusions made here are contained in the article or Supporting Information. Any other data is available at request from the authors.

## Supporting information

This article contains supporting information (104-137).

## Acknowledgements

The computational resources were provided by The University of Chicago Research Computing Center (RCC).

## Author contributions

Z. Y. and G. A. V. designed research. Z. Y., D. T., and Z. W. performed simulations. Z. Y. and J. W. analyzed data. Z. Y. and G. A. V. wrote the manuscript.

## Funding and additional information

This research was supported in part by the National Institute of General Medical Sciences (NIGMS) of the National Institutes of Health (NIH) through grant R01GM053148 to G.A.V. The content is solely the responsibility of the authors and does not necessarily represent the official views of the National Institutes of Health. The research was also supported in part by the Office of Naval Research (Award N00014-21-1-2157 to G.A.V.).

## Abbreviations–The abbreviations used are

BM2: influenza B M2 proton channel
AM2: influenza A M2 proton channel
NT: N-terminal
CT: C-terminal
FPS: fixed-protonation-state molecular dynamics
CpHMD: continuous constant-pH molecular dynamics
PT: proton transport
MS-RMD: multiscale reactive molecular dynamics
QM/MM: quantum mechanics/molecular mechanics
HBond: hydrogen-bond
GB: generalized-Born
WT: wildtype
ssNMR: solid-state NMR
POPC: 1-palmitoyl-2-oleoyl-*sn*-glycero-3-phosphocholine
VM+: virus-mimetic
RMSD: root-mean-squared deviation
POPE: 1-palmitoyl-2-oleoyl-*sn*-glycero-3-phosphoethanolamine

