## Supporting Information for "Activation of the influenza B M2 proton channel (BM2)"

### Supporting Information for
### Activation of the influenza B M2 proton channel (BM2)

Zhi Yue, Jiangbo Wu, Da Teng, Zhi Wang,† and Gregory A. Voth*

*Department of Chemistry, Chicago Center for Theoretical Chemistry, James Franck Institute, and Institute for Biophysical Dynamics, The University of Chicago, Chicago, Illinois 60637, USA*

†Present address: HAKUHODO Technologies Inc., Minato, Tokyo, 107-6320, Japan.

### 1 Survey of BM2 PDB structures

**Table S1. Comparison of published structures of BM2**

| PDB | Method[a] | Resolution (Å) | pH[b] | T (K) | Lipid[c] | Mutation[d] | Conformation | RMSD[e] (Å) | Ref. |
|---|---|---|---|---|---|---|---|---|---|
| 2KIX | NMR | - | 7.5 | 305 | DHPC | C11S | closed | 14.7 | Chou et al. (1) |
| 2KJ1[f] | NMR | - | 6.8 | 305 | LMPG | - | - | - | |
| 6PVR | ssNMR | - | 7.5 | 280–290 | POPE | - | closed | - | Hong et al. (2) |
| 6PVT | | | 4.5 | | | - | open | 4.1 | |

[a] ssNMR: solid-state NMR; NMR: solution NMR. [b] pH where NMR experiments were conducted. [c] DHPC (detergent): dihexanoylphosphocholine; LMPG (detergent): *lyso*-myristoylphosphatidylglycerol; POPE: 1-palmitoyl-2-oleoyl-*sn*-glycero-3-phosphoethanolamine. Note that POPE lipids at 280–290 K are in a gel phase (3-6). [d] Wild-type sequence shown in Figure 1C. [e] Backbone RMSD of residues 1–33 (resolved in all PDBs) with respect to the reference structure. [f] Cytoplasmic domain, residues 43–103.

**Table S2. Perimeter of the pore-lining residues in the ssNMR BM2 structures**

| Pore-lining residue | Quadrilateral perimeter of $C_\alpha$ atoms (Å) | | |
|---|---|---|---|
| | 6PVR (closed)[a] | 6PVT (open)[a] | Change upon activation[b] |
| Phe5 | 31.5 ± 1.7 | 59.9 ± 2.2 | 28.3 ± 2.8 |
| Leu8 | 38.2 ± 2.2 | 52.2 ± 2.9 | 14.0 ± 3.6 |
| Ser12 | 36.7 ± 1.5 | 46.3 ± 3.0 | 9.6 ± 3.4 |
| Ser16 | 41.3 ± 1.1 | 47.5 ± 3.1 | 6.2 ± 3.2 |
| His19 | 42.4 ± 0.6 | 48.6 ± 1.5 | 6.2 ± 1.6 |
| Trp23 | 44.5 ± 1.2 | 45.3 ± 1.6 | 0.8 ± 2.0 |
| His27 | 51.3 ± 2.8 | 45.7 ± 3.8 | –5.6 ± 4.7 |

[a] Reported as mean ± standard deviation computed from all NMR models. [b] Errors computed from error propagation.

### 2 Simulation Protocols

#### 2.1 System preparation

The first models of closed (PDB ID: 6PVR) and open (PDB ID: 6PVT) ssNMR structures (2) were re-oriented with respect to the lipid bilayer using the Positioning of Proteins in Membranes (PPM) 2.0 web server (https://opm.phar.umich.edu/ppm_server) (7) for the starting BM2 configurations.

Following the CHARMM-GUI protocol (8, 9), systems were built using CHARMM (10) (version c42b2). BM2 residues 1–33 were included, with the N-terminal (NT) and C-terminal (CT) respectively acetylated and amidated. Hydrogen atoms were generated using the HBUILD facility (11). To relax the positions of hydrogen atoms, energy minimization was performed using steepest descent (12) (SD) for 50 steps followed by adopted basis Newton-Raphson (13) (ABNR) for 50 steps, with heavy atoms fixed.

The minimization was done in implicit solvent represented by the generalized-Born (GB) model GBSW (14) (GB model with a smoothed switching function) with optimized input radii (15). Pore waters (PWAT) were added to fill protein cavities. Consistent with previous AM2 simulations (16-20), proteins were embedded into a lipid bilayer of 163 POPC (1-palmitoyl-2-oleoyl-*sn*-glycero-3-phosphocholine) molecules. A water layer (BWAT) of about 14–15 Å thick was added above and below the lipid bilayer to ensure a similar number of atoms in all systems. $Cl^-$ ions (BION) were added to the bulk water to neutralize the systems. No extra ions were added because the histidine p$K_a$s were determined under an ionic strength of ~20 mM (21-23). Bulk water molecules within 2.8 Å of existing heavy atoms were removed. The dimensions of the final systems are about 78Å × 78Å × 85Å. CHARMM structures were converted to the GROMACS format using GROMACS (24) (version 5.1.5) as needed. The details of the systems (Closed_POPC and Open_POPC) are listed in **Table S3**.

**Table S3. Details of the simulation systems**

| System | Charge states[a] | Number of atoms/molecules[b] | | | | | |
|---|---|---|---|---|---|---|---|
| | | Protein[c] | PWAT | Lipid[d] | BWAT | BION | Total |
| Open_POPC | $H19^{+3}/H27^{+4}$ | 2247 | 37 | 163 POPC | 9025 | 11 $Cl^-$ | 51286 |
| Open_VM+ | | | 30 | 41 POPC + 41 POPE + 41 PSM + 40 CHL1 | 9030 | | 48224 |
| Open_POPE | | | 40 | 163 POPE | 9030 | | 49843 |
| Closed_POPC | $H19^{+0}/H27^{+1}$ | 2241 | 20 | 163 POPC | 9048 | 5 $Cl^-$ | 51292 |
| Closed_VM+ | | | 17 | 41 POPC + 41 POPE + 41 PSM + 40 CHL1 | 9046 | | 48221 |
| Closed_POPE | | | 18 | 163 POPE | 9046 | | 49813 |
| H27A_POPC[e] | $H19^{+0}$ | 2224 | 20 | 163 POPC | 9048 | 5 $Cl^-$ | 51275 |

[a] E3s were deprotonated while K32s and R33s were protonated. [b] PWAT: pore waters introduced to fill protein cavities; BWAT: bulk waters; BION: bulk ions. [c] A wild-type tetramer consists of 4 GLU, 8 HIS, 4 LYS, and 4 ARG residues. [d] POPC: 1-palmitoyl-2-oleoyl-*sn*-glycero-3-phosphocholine; POPE: 1-palmitoyl-2-oleoyl-*sn*-glycero-3-phosphoethanolamine; PSM: N-palmitoyl-D-erythro-sphingosylphosphorylcholine; CHL1: cholesterol. [e] Built from the equilibrated Closed_POPC system.

Note that the histidine p$K_a$s and the NMR structures were determined in virus-mimetic (VM+) (21-23) and 1-palmitoyl-2-oleoyl-*sn*-glycero-3-phosphoethanolamine (POPE) membranes (2), respectively. The VM+ membrane comprises POPC:POPE:sphingomyelin:cholesterol with a molar ratio of 1:1:1:1 (21-23). As per Luo et al. (25), the sphingomyelin used was egg sphingomyelin obtained from Avanti® Polar Lipids, which consists of 86% N-palmitoyl-D-erythro-sphingosylphosphorylcholine (PSM) (26). To evaluate the lipid impact on BM2 dynamics, four additional systems (Closed_VM+, Open_VM+, Closed_POPE, and Open_POPE) were built (**Table S3**).

CpHMD simulations began with the final snapshot of bilayer equilibration (refer to section 1.3). Note that charge states of ionizable residues in fixed-protonation-state (FPS) simulations don't affect CpHMD simulations because protein heavy atoms were restrained in FPS equilibration.

The mutant system (H27A_POPC, **Table S3**) was derived from the equilibrated closed wildtype (WT) system (Closed_POPC). The system's net charge was not adjusted because the fluctuating net charge is dynamically offset in the CpHMD simulation (refer to section 1.2).

#### 2.2 Simulation settings

Simulations were performed using CHARMM (10) (version c47b1) and GPU-accelerated GROMACS (27, 28) (version 2019.3). CHARMM employed the hybrid-solvent CpHMD and pH-based replica exchange (pH-REX) methods (29) via PHMD and REPDSTR modules, respectively. Proteins were modeled by the CHARMM22 (30) force field with CMAP (grid-based energy correction term) correction (31, 32), while lipids were represented by the CHARMM36 model (33-35), waters by the CHARMM-modified (36) TIP3P model (37), and ions by the Beglov and Roux Lennard-Jones (LJ) model (38) with NBFIX terms (39, 40). CHARMM used the Verlet leap-frog integrator (41) to propagate spatial coordinates. GROMACS used the velocity Verlet (42) and Verlet leap-frog integrators for the NVT (constant temperature and volume) and NPT (constant temperature and pressure) ensembles, respectively. Bonds involving hydrogen atoms were restrained using SHAKE (43) in CHARMM or LINCS (44) in GROMACS to allow for a 2-fs step length. LJ interactions were force-switched (45) from 8 to 12 Å. Coulombic interactions were computed by the particle mesh Ewald (PME) method (46, 47) with a real-space cutoff of 12 Å, an approximate 1-Å grid spacing, and sixth-order and fourth-order interpolations in CHARMM and GROMACS, respectively. The non-bonded neighbor list was updated heuristically in CHARMM or every 20 fs in GROMACS. Simulations were performed under periodic boundary conditions (PBC) at 308.15 K maintained by the Nosé–Hoover thermostat (48, 49). In NPT simulations, a pressure of 1 atm in CHARMM and of 1 bar in GROMACS was semiisotropically maintained by the Langevin piston pressure-coupling algorithm (50) and the Parrinello-Rahman barostats (51), respectively. Masses of thermal and pressure pistons were set to 5000 kcal•ps$^2$/mol and 2500 amu (atomic mass unit) respectively in CHARMM. Time constants for coupling were set to 1 and 5 ps respectively for thermostat and barostat in GROMACS, following the protocol optimized by Lee et al. (52). CHARMM and GROMACS simulations collected data every 1 and 10 ps, respectively.

CpHMD propagates fictitious λ particles of 10 amu in mass associated with each ionizable residue using the Langevin algorithm (53) with a collision frequency of 5 ps$^{-1}$. To offset the net charge fluctuations resulting from the dynamic protonation of ionizable residues in CpHMD, charge neutrality is maintained by compensating the net charge using background plasma in the PME method (46, 47). In hybrid-solvent CpHMD, explicit solvent and lipid molecules are utilized to propagate spatial coordinates, while implicit solvent and membrane modeled by the membrane-enabled (54) GBSW (14) model with optimized GB input radii (15) are utilized to compute electrostatic hydration forces on titration coordinates (29, 55). The implicit solvent and membrane are represented by a continuous medium with a dielectric constant of 80 and an infinite slab with a dielectric constant ($\varepsilon_m$) of 1, respectively, with the salt screening effect approximated by a Debye–Hückel term in GB (56). To exclude the impact of implicit membrane on the protonation of buried protein ionizable residues, a high-dielectric exclusion cylinder placed along the membrane normal (Z-axis). In the BM2 simulations, an implicit membrane of 33 Å in thickness (the apparent hydrophobic thickness of the POPC bilayer (57)) was applied. $\varepsilon_m$ was switched from 1 to 80 within 2.5 Å from each implicit membrane surface. GB calculations were done with 38 angular integration points and 50 radial integration points, which have been shown to reproduce the electrostatic solvation energy obtained from Poisson–Boltzmann calculations with $\varepsilon_m$ = 1 (54). Consistent with the AM2 CpHMD simulation by Chen et al. (55), an exclusion cylinder with a radius of 15 Å was employed. Since only counter ions were added, an ionic strength of 0 mM was used in the Debye–Hückel term. Titration coordinates were updated every 20 fs to relax water molecules around ionizable residues (29). With pH-REX activated, neighboring pH replicas attempted to exchange every 1 ps, with acceptance determined using the Metropolis criterion (58).

Undocumented options defaulted to the standard settings in CHARMM or GROMACS.

#### 2.3 Simulation protocol

Adapted from previous studies (59, 60), simulations were performed in three stages (**Table S4**).

Stage 1. Initial equilibration employed CHARMM to eliminate bad contacts, reposition components appropriately, and stabilize system dimensions. After multiple energy minimizations using the SD and

ABNR methods, the system underwent gradual relaxation in six steps by applying various restraints to protein, lipids, waters, and ions.

Stage 2. Bilayer equilibration ensured POPC bilayer equilibrium by equilibrating the last configuration from stage 1 using GROMACS. After minimization using the SD method with protein and lipid heavy atoms restrained until the maximum force dropped below 1000 kJ/mol•nm, the system underwent NVT equilibration for 1 ns followed by NPT equilibration for 100 ns, with protein restraints maintained. The FPS production run lasted for 2.9 µs without restraints.

Stage 3. CpHMD run used the last configuration from stage 2.2 to initiate a membrane-enabled (55) hybrid-solvent (29) CpHMD simulation in CHARMM. To enable titration, doubly protonated HIS residues were used, and dummy hydrogen atoms were added to GLU carboxylates at *syn* positions (61). Dummy hydrogen positions were relaxed through initial minimization using the SD and ABNR methods, with protein heavy atoms fixed and lipid heavy atoms restrained. Subsequent equilibrations totaling 1 ns were conducted at pH 7.5 (where the ssNMR structure was determined) with gradual release of protein positional restraints. Starting from the fourth equilibration, a cylindrical restraint was applied to the center of mass (COM) of protein heavy atoms to prevent lateral diffusion. In the pH-REX production run, 15 replicas spanning pH 2.5 to 9.5 with an interval of 0.5 lasted for 100 ns per replica, accumulating a 1.5-µs total sampling time. Throughout the CpHMD run, bulk ions were restrained by a planar potential to prevent their penetration into the bilayer region.

**Table S4. Details of the simulation protocol (POPC and VM+ lipids)**

| Stage | Ensemble | Step length (fs) | Length (ns) | Force constant in the restraint potential[a] | | | | |
|---|---|---|---|---|---|---|---|---|
| 1. Initial equilibration | | | | BB | SC+PWAT | Lipid[b] | BWAT[c] | BION[c] |
| 1 | NVT[d] | 1 | 0.1 | 10.0 | 5.0 | 2.5 | 2.5 | 2.5 |
| 2 | NVT[d] | 2 | 0.1 | 5.0 | 5.0 | 2.5 | 2.5 | 2.5 |
| 3 | NPT | 2 | 0.2 | 5.0 | 5.0 | 1.0 | 1.0 | 1.0 |
| 4 | NPT | 2 | 0.2 | 2.5 | 2.5 | 0.5 | 0.5 | 1.0 |
| 5 | NPT | 2 | 0.2 | 1.5 | 1.5 | 0.1 | 0.1 | 1.0 |
| 6 | NPT | 2 | 1.2 | 1.0 | 1.0 | - | - | 1.0 |
| 2. Bilayer equilibration & FPS production run | | | | BB | SC | Lipid | Water | BION |
| 1 | NVT | 2 | 1.0 | 1.0 | 1.0 | - | - | - |
| 2 | NPT | 2 | 100 | 1.0 | 1.0 | - | - | - |
| 3 | NPT | 2 | 2900 | - | - | - | - | - |
| 3. CpHMD run | | | | BB | SC | Lipid | Water | BION[f] |
| 1 | NVT[d] | 1 | 0.1 | 1.0 | 1.0 | - | - | 1.0 |
| 2 | NPT | 2 | 0.2 | 1.0 | 1.0 | - | - | 1.0 |
| 3 | NPT | 2 | 0.2 | 0.5 | 0.5 | - | - | 1.0 |
| 4 | NPT[e] | 2 | 0.5 | - | - | - | - | 1.0 |
| 5 | NPT[e] | 2 | 100 per replica | - | - | - | - | 1.0 |

[a] Harmonic positional and other restraints were respectively applied using the *cons* command and the MMFP facility in CHARMM (10). BB and SC stand for protein backbone and side-chain heavy atoms, respectively. PWAT, BWAT and BION represent pore water, bulk water molecules and ions, respectively. [b] Planar potentials restrained the tails of lipid molecules near the bilayer center (–5 Å < Z < 5 Å) and headgroups near the bilayer surface (Z = ±19 Å for POPC and POPE; Z = ±21 Å for PSM; Z = ±18 Å for CHL1). [c] A planar potential was applied to the oxygen atoms of bulk waters (–11 Å < Z < 11 Å) or bulk ions (–15 Å < Z < 15 Å) to exclude them from the hydrophobic region of the bilayer. [d] The Langevin dynamics (53) was employed. [e] A cylindrical potential of 1.0 kcal/mol•Å$^2$ was added to the protein center of mass to prevent lateral diffusion. [f] Bulk ions were prevented from entering the bilayer region (–16.5 Å < Z < 16.5 Å) by a planar potential (62).

Systems Closed_POPE and Open_POPE were designed to study BM2 behaviors under experimental conditions (POPE membranes at 280–290 K). Starting from the closed or open configuration, the systems were first equilibrated at 308.15 K with protein heavy atoms restrained to obtain a fluid-phase POPE bilayer (stages 1–2 in **Table S5**). Then, the temperature was reduced to 280.15 K and the simulations ran for 1.5 μs with protein restraints maintained to induce transition to the gel-phase (stage 3.2 in **Table S5**). The simulations were then continued for 5.0 μs with protein restraints removed to relax the protein in the gel-phase (stage 3.3 in **Table S5**).

**Table S5. Details of the simulation protocol (POPE lipids)**

| Stage | Ensemble | Step length (fs) | Length (ns) | Force constant in the restraint potential[a] (kcal/mol•Å$^2$) | | | | |
|---|---|---|---|---|---|---|---|---|
| 1. Initial equilibration | | (308.15 K) | | BB | SC+PWAT | Lipid[b] | BWAT[c] | BION[c] |
| 1 | NVT[d] | 1 | 0.1 | 10.0 | 5.0 | 2.5 | 2.5 | 2.5 |
| 2 | NVT[d] | 2 | 0.1 | 5.0 | 5.0 | 2.5 | 2.5 | 2.5 |
| 3 | NPT | 2 | 0.2 | 5.0 | 5.0 | 1.0 | 1.0 | 1.0 |
| 4 | NPT | 2 | 0.2 | 2.5 | 2.5 | 0.5 | 0.5 | 1.0 |
| 5 | NPT | 2 | 0.2 | 1.5 | 1.5 | 0.1 | 0.1 | 1.0 |
| 6 | NPT | 2 | 1.2 | 1.0 | 1.0 | - | - | 1.0 |
| 2. Fluid-phase equilibration | | (308.15 K) | | BB | SC | Lipid | Water | BION |
| 1 | NVT | 2 | 1.0 | 1.0 | 1.0 | - | - | - |
| 2 | NPT | 2 | 100 | 1.0 | 1.0 | - | - | - |
| 3. Gel-phase equilibration | | (280.15 K) | | BB | SC | Lipid | Water | BION |
| 1 | NVT | 2 | 1.0 | 1.0 | 1.0 | - | - | - |
| 2 | NPT | 2 | 1500 | 1.0 | 1.0 | - | - | - |
| 3 | NPT | 2 | 5000 | - | - | - | - | - |

[a] Harmonic positional and other restraints were respectively applied using the *cons* command and the MMFP facility in CHARMM (10). BB and SC stand for protein backbone and side-chain heavy atoms, respectively. PWAT, BWAT and BION represent pore water, bulk water molecules and ions, respectively. [b] Planar potentials restrained the tails of lipid molecules near the bilayer center (–5 Å < Z < 5 Å) and headgroups near the bilayer surface (Z = ±19 Å for POPE). [c] A planar potential was applied to the oxygen atoms of bulk waters (–11 Å < Z < 11 Å) or bulk ions (–15 Å < Z < 15 Å) to exclude them from the hydrophobic region of the bilayer. [d] The Langevin dynamics (53) was employed.

### 3 Analysis Protocols

#### 3.1 p$K_a$ calculation

To calculate the microscopic p$K_a$ of an ionizable residue, the unprotonated fraction $S^{\mathrm{unprot}}$ was first computed by counting the populations of protonated (defined as those with $\lambda \leq 0.1$) and deprotonated ($\lambda \geq 0.9$) states at a specific pH. The p$K_a$ was then computed by fitting $S^{\mathrm{unprot}}$ *vs.* pH to the Hill equation (63),

$$S^{\mathrm{unprot}} = \frac{1}{1+10^{n(\mathrm{p}K_a - \mathrm{pH})}}, \quad \text{(Eq. S1)}$$

where the Hill coefficient $n$ describes the steepness of the transition region in a titration curve.

The stepwise protonation of a histidine tetrad involves four equilibria. Following Chen et al. (55), the $S^{\mathrm{unprot}}$s were calculated as follows.

$$S_{0\text{-}1} = \frac{[\mathrm{His}_4^0]}{[\mathrm{His}_4^0] + [\mathrm{His}_4\mathrm{H}^+]} \quad \text{(Eq. S2)}$$

$$S_{1\text{-}2} = \frac{[\mathrm{His_4H^+}]}{[\mathrm{His_4H^+}] + [\mathrm{His_4H_2^{2+}}]}$$

$$S_{2\text{-}3} = \frac{[\mathrm{His_4H_2^{2+}}]}{[\mathrm{His_4H_2^{2+}}] + [\mathrm{His_4H_3^{3+}}]}$$

$$S_{3\text{-}4} = \frac{[\mathrm{His_4H_3^{3+}}]}{[\mathrm{His_4H_3^{3+}}] + [\mathrm{His_4H_4^{4+}}]}$$

Stepwise p$K_1$, p$K_2$, p$K_3$, and p$K_4$ were then computed by respectively fitting $S_{3\text{-}4}$, $S_{2\text{-}3}$, $S_{1\text{-}2}$, and $S_{0\text{-}1}$ to Eq. S1.

#### 3.2 Other analyses

The CpHMD trajectories were analyzed using CHARMM (version c42b2). RMSD was calculated with the *coor rms* command by least-square fitting to the backbone atoms (N, $C_\alpha$, C) using *coor orient rms*. The OPM structure of 6PVR served as the reference. Atomic distances were calculated using the *quick* command, while the minimal distance between two groups of atoms was computed using *coor mind*. Principal axes were computed using the *coor iner* command.

The GROMACS trajectories were analyzed using GROMACS (version 5.1.5). RMSD was calculated with the *gmx rms* command by least-square fitting to the backbone atoms (N, $C_\alpha$, C). The OPM structure of 6PVR served as the reference. Atomic distances were calculated using the *gmx mindist* command.

The distribution of water molecules along the channel ($N_{Water}$) was defined as the number of water oxygen atoms in 2-Å slices within a radius of 15.0 Å centered along the Z-axis and computed using the MDAnalysis (64) script developed by Gelenter et al. (65). The COM of a residue was computed for side-chain heavy atoms and projected onto the channel axis using MDAnalysis (64). The COM of four Ala17 $C_\alpha$ atoms was used as the origin for the projection.

#### 3.3 Statistical analysis

Unless otherwise specified, the initial 40 ns of a CpHMD run were excluded from analyses. All reported errors were standard deviations calculated from block analysis, with 40–100 ns divided into three 20-ns blocks. For FPS runs with POPC and VM+ membranes, the last 2000 ns were used for analyses based on the time series of RMSD (**Figure S11**). For FPS runs with POPE membranes, the last 3000 ns were used for analyses based on the time series of RMSD (**Figure S18**).

#### 3.4 PDB survey

All BM2 structures deposited in the PDB bank (**Table S1**) were analyzed. The OPM structure of 6PVR was chosen as the reference, and other PDB structures were aligned to it by least-square fitting the backbone atoms. Analyses were conducted using in-house Tcl scripts for Visual Molecular Dynamics (66) (VMD, version 1.9.4a57).

#### 3.5 Figure generation

Structural drawings were generated using VMD (66) (version 1.9.4a57), and data plots were created using Grace (67) (version 5.1.25).

### 4 Supplementary Figures and Tables

#### 4.1 Figures and tables discussed in the main text

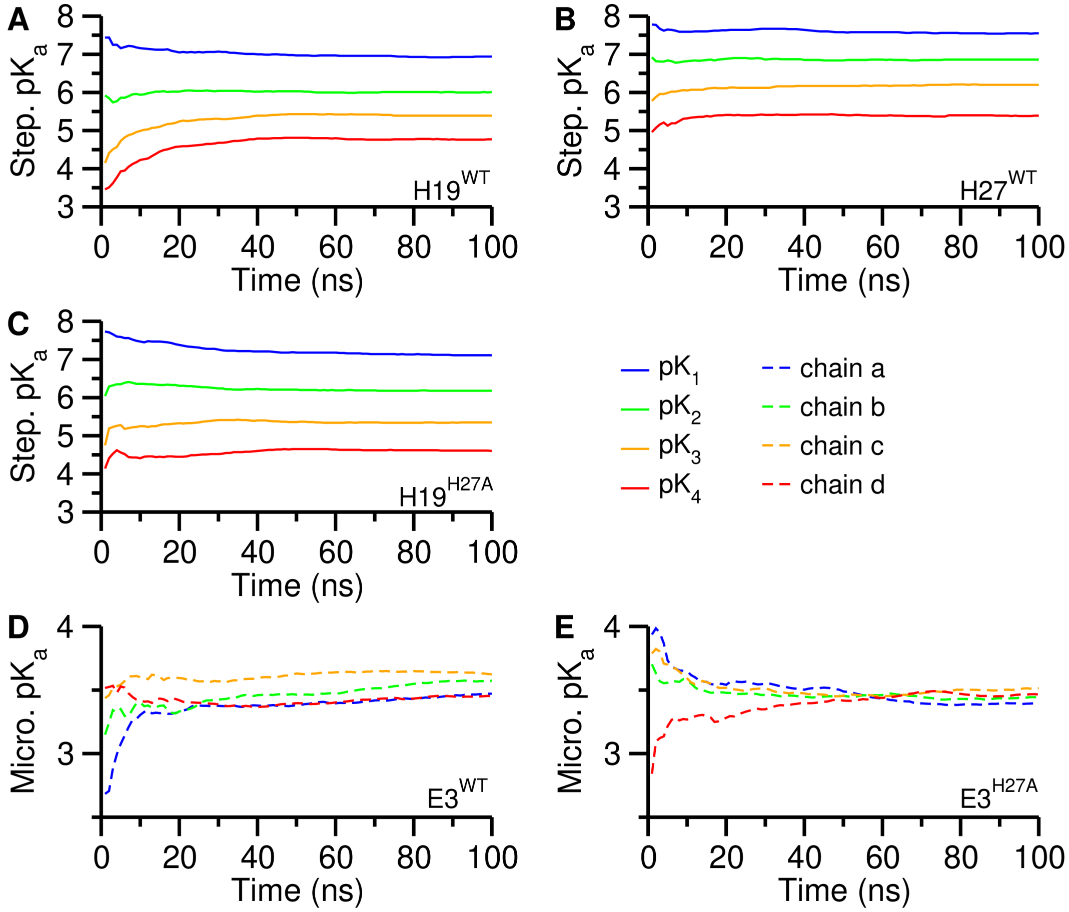


**Figure S1. Convergence of HIS stepwise (A–C) and GLU microscopic (D–E) p$K_a$s in the CpHMD simulations.** p$K_a$s were calculated cumulatively as a function of simulation time. Superscripts "WT" and "H27A" in legends stand for the wild-type and H27A mutant, respectively.

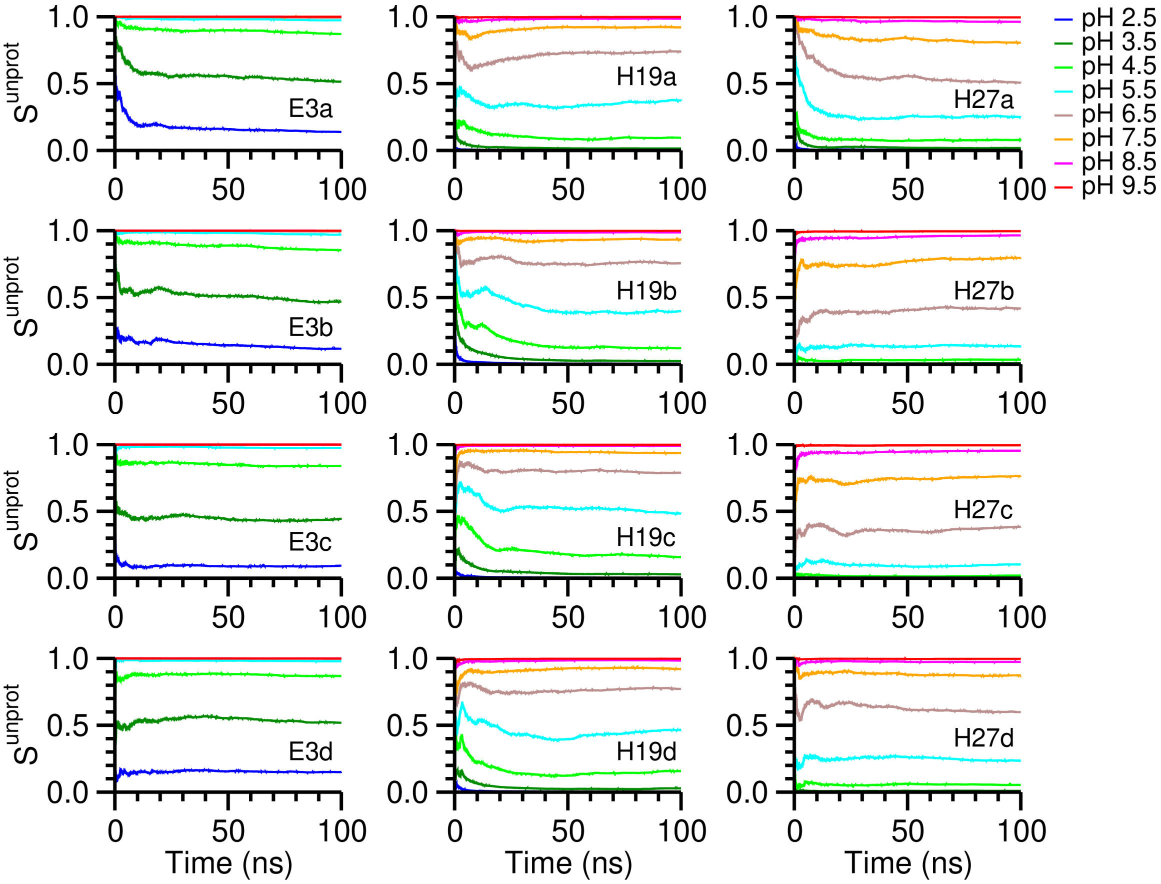


**Figure S2. Convergence of $S^{unprot}$s in the WT CpHMD simulation.** $S^{unprot}$s of Glu3, His19, and His27 were calculated cumulatively as a function of simulation time. For clarity, only half-integer pH values are displayed.

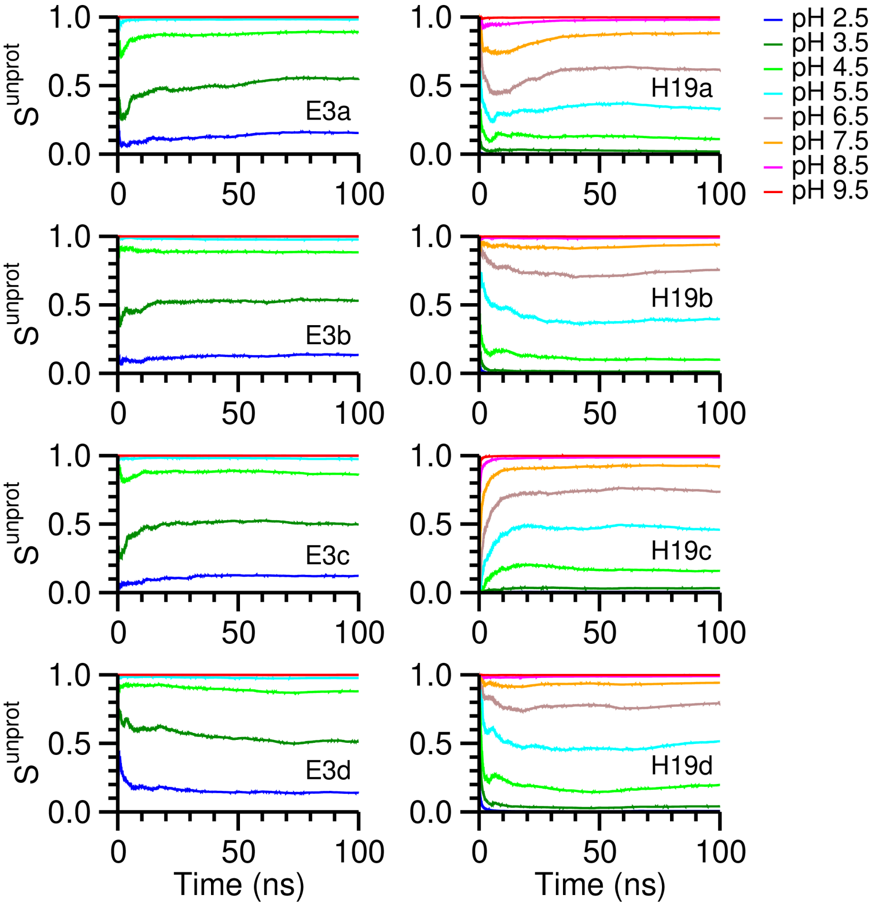


**Figure S3. Convergence of $S^{unprot}$s in the H27A CpHMD simulation.** $S^{unprot}$s of Glu3 and His19 were calculated cumulatively as a function of simulation time. For clarity, only half-integer pH values are displayed.

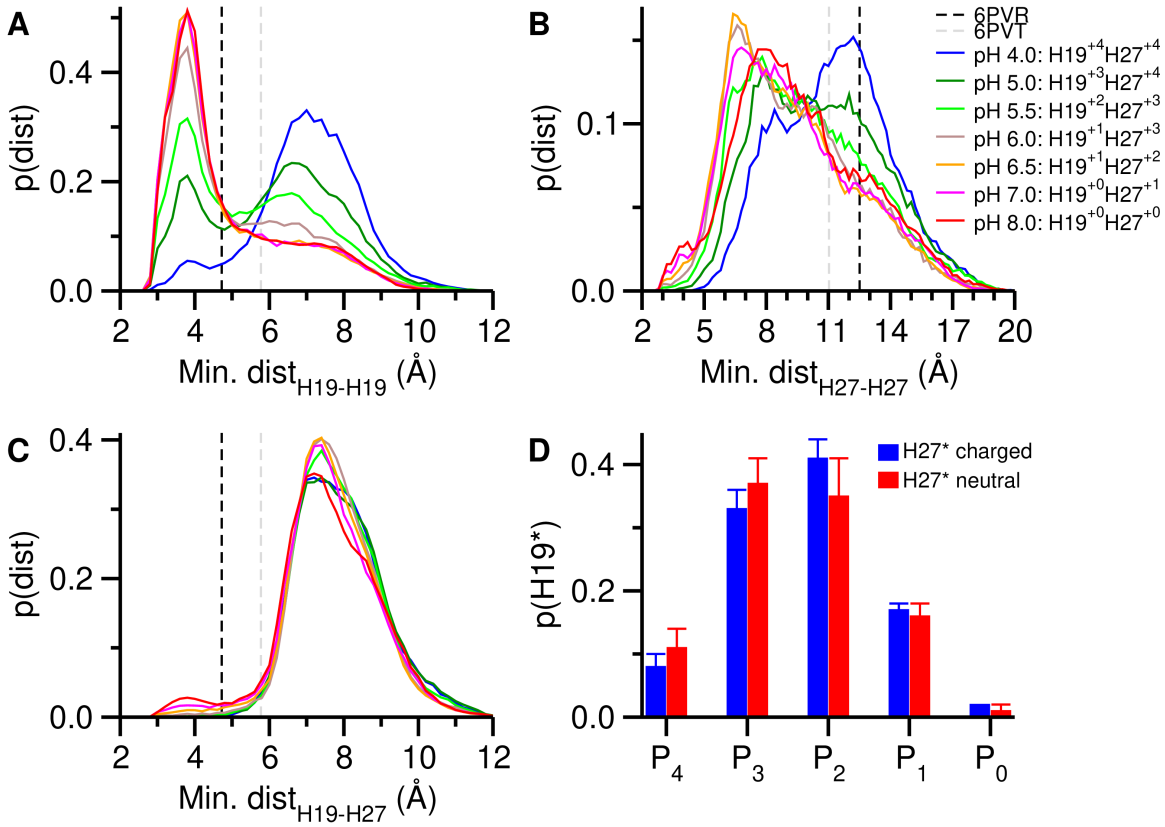


**Figure S4. Impact of His27 protonation states on that of His19.** Probability distributions of the minimum distance between H19 and H19 (A), H27 and H27 (B), and H19 and H27 (C) in the WT CpHMD simulation. The minimum distances were measured using side-chain nitrogen atoms ($N^{\sigma 1}$ and $N^{\varepsilon 2}$). The vertical black and grey dashed lines represent the closed (PDB ID: 6PVR) and open (PDB ID: 6PVT) ssNMR structures, respectively. (D) Probability distribution of various protonation states of His19 ($P_0/P_1/P_2/P_3/P_4$) at pH 5.5 when the closest His27 (H27*) is charged (blue bars) and neutral (red bars).

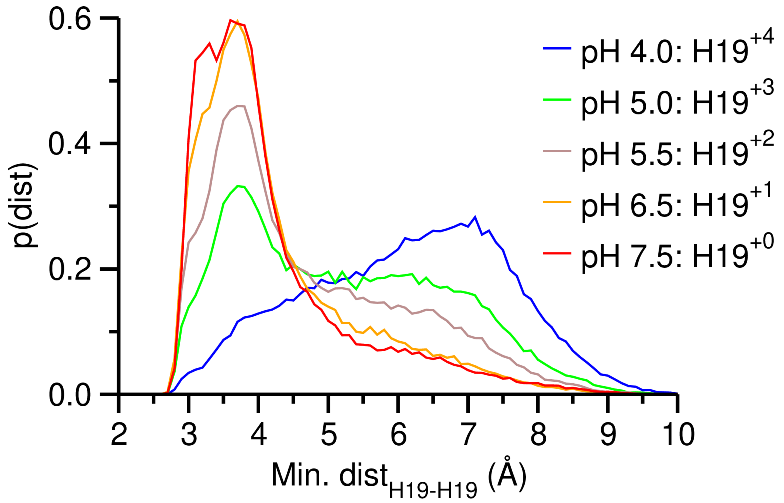


**Figure S5. Probability distributions of the minimum distance between His19s in the H27A mutant simulation.** The minimum distances were measured using side-chain nitrogen atoms ($N^{\sigma 1}$ and $N^{\varepsilon 2}$).

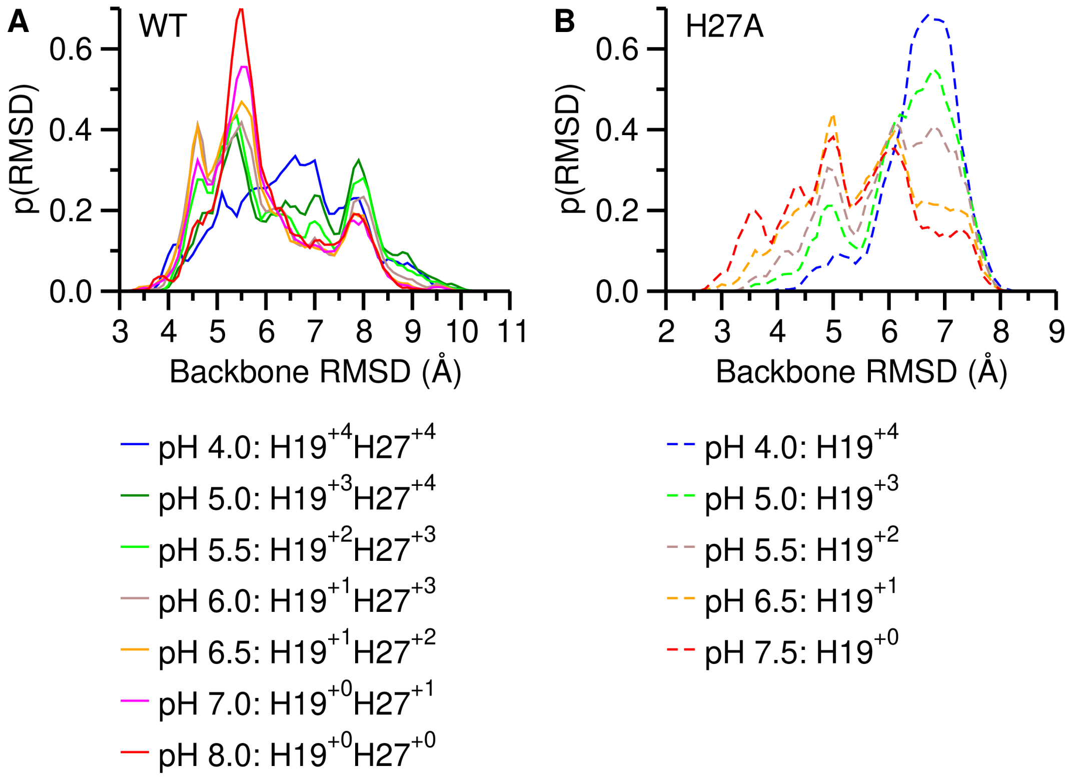


**Figure S6. Probability distribution of backbone RMSDs in the WT (A) and H27A mutant (B) CpHMD simulations.**

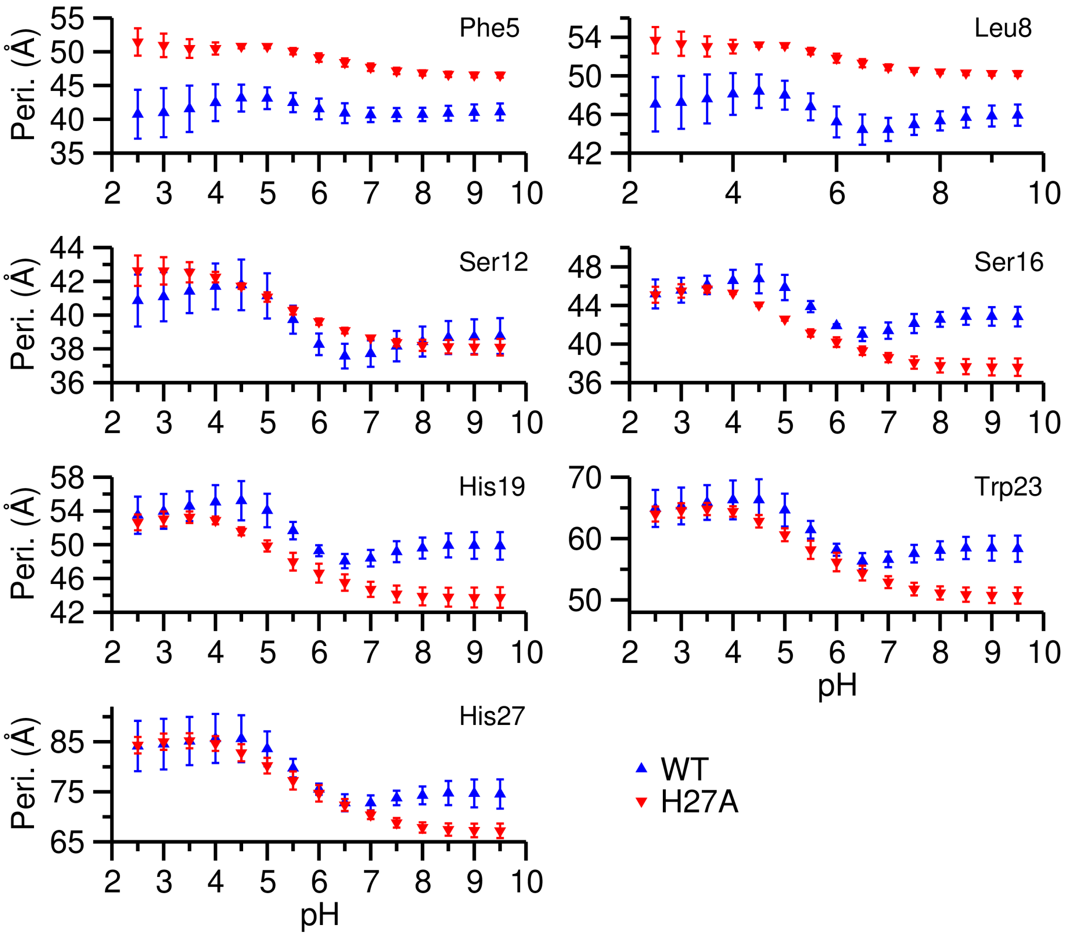


**Figure S7. Comparison of perimeters of the WT and H27A mutant.** Means over the simulation time were plotted versus pH values and errors were standard deviations estimated from block analysis.

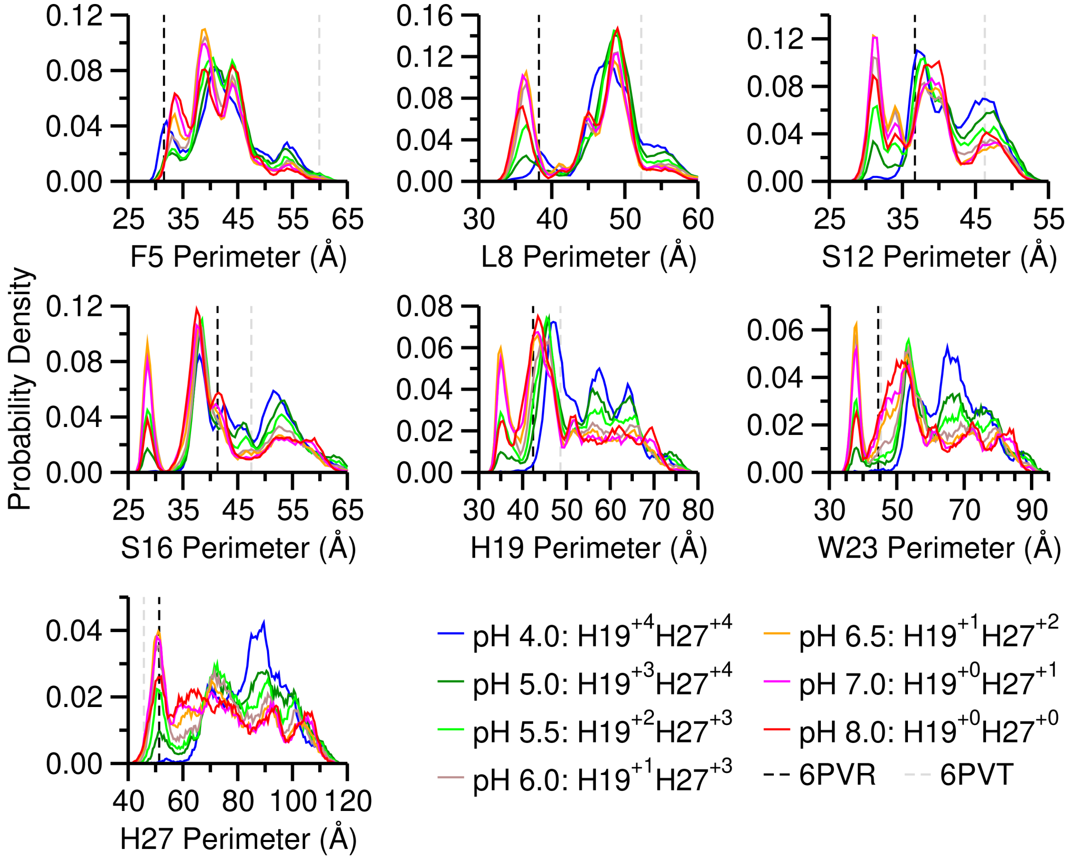


**Figure S8. Probability distribution of perimeters of the pore-lining residues in the WT CpHMD simulation.** A perimeter was computed for the quadrilateral connecting the $C_\alpha$ atoms of four symmetric residues. The vertical black and grey lines represent the closed (PDB ID: 6PVR) and open (PDB ID: 6PVT) ssNMR structures, respectively.

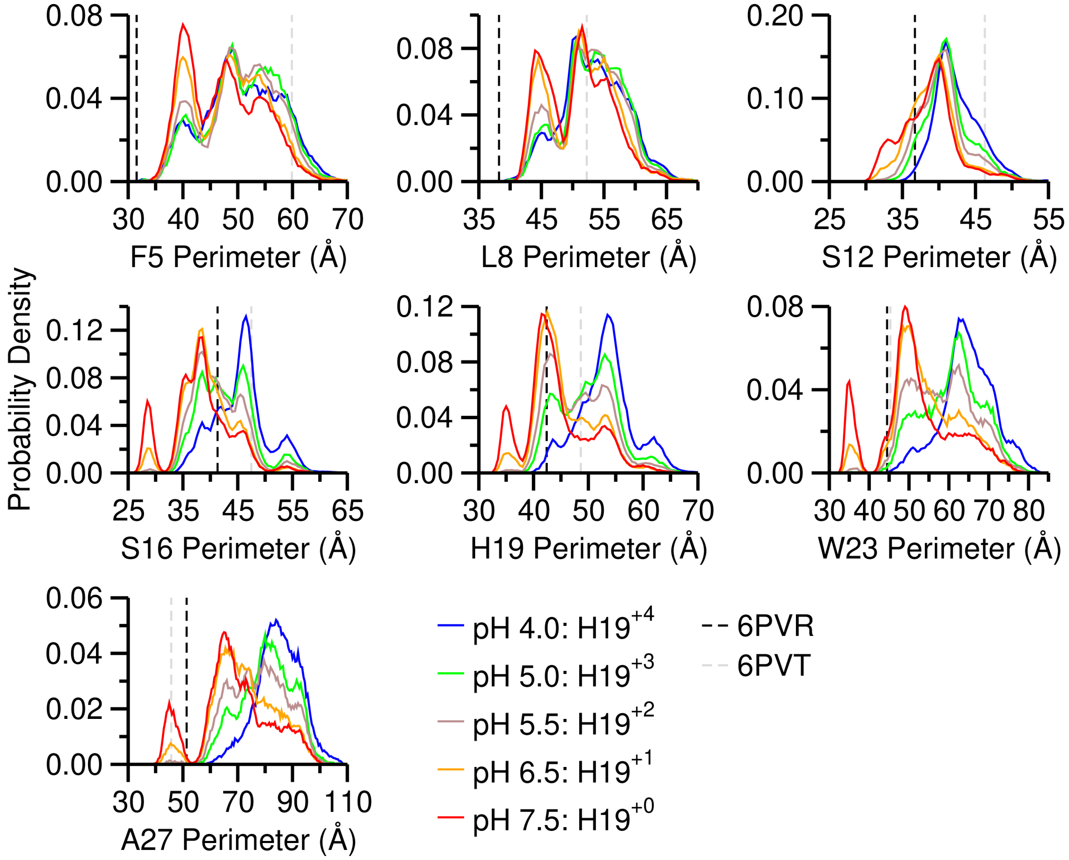


**Figure S9. Probability distribution of perimeters of the pore-lining residues in the H27A mutant CpHMD simulation.** A perimeter was computed for the quadrilateral connecting the $C_\alpha$ atoms of four symmetric residues. The vertical black and grey lines represent the closed (PDB ID: 6PVR) and open (PDB ID: 6PVT) ssNMR structures, respectively.

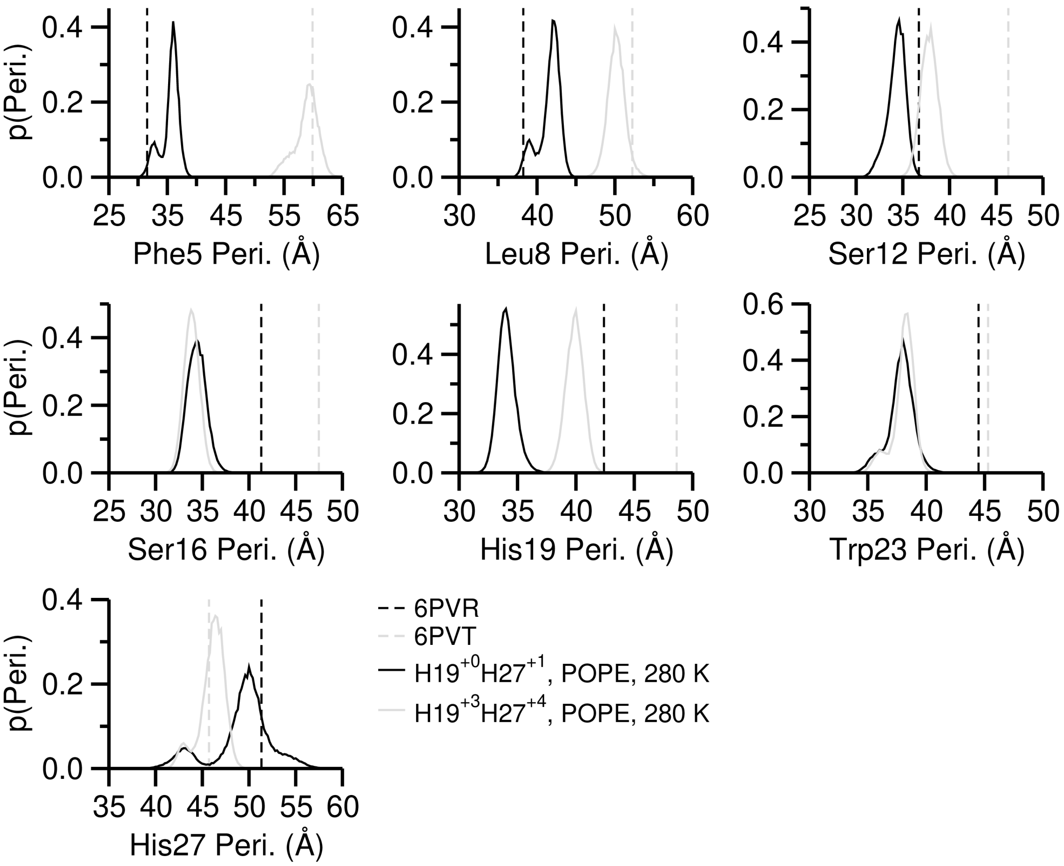


**Figure S10. Perimeters of the pore-lining residues in gel-phase bilayers.** Probability distributions of perimeters of the pore-lining residues in the closed (black solid lines) and open (grey solid lines) FPS simulations with POPE bilayers at 280.15 K. A perimeter was computed for the quadrilateral connecting the $C_\alpha$ atoms of four symmetric residues. The vertical black and grey dashed lines represent the closed (PDB ID: 6PVR) and open (PDB ID: 6PVT) ssNMR structures, respectively.

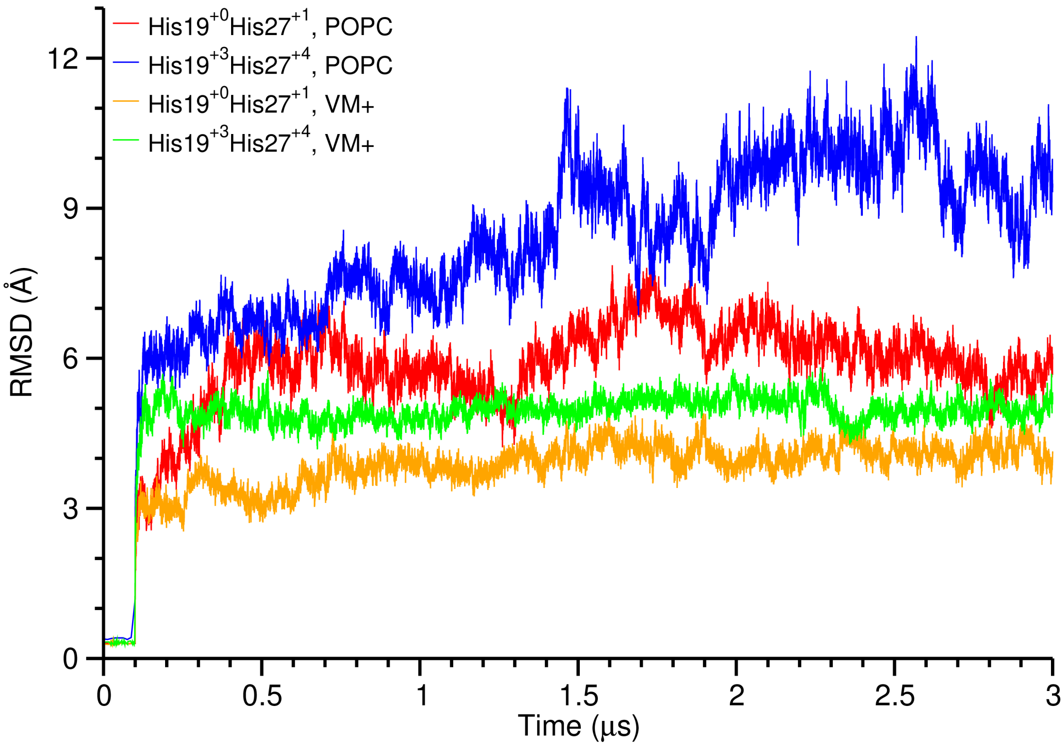


**Figure S11. Backbone RMSDs of the WT BM2 in fluid-phase bilayers.** Time series of RMSD in the closed FPS simulation with a POPC bilayer (red), the open FPS simulation with a POPC bilayer (blue), the closed FPS simulation with a VM+ bilayer (orange), and the open FPS simulation with a VM+ bilayers (green) at 308.15 K. Sampled every 100 ps.

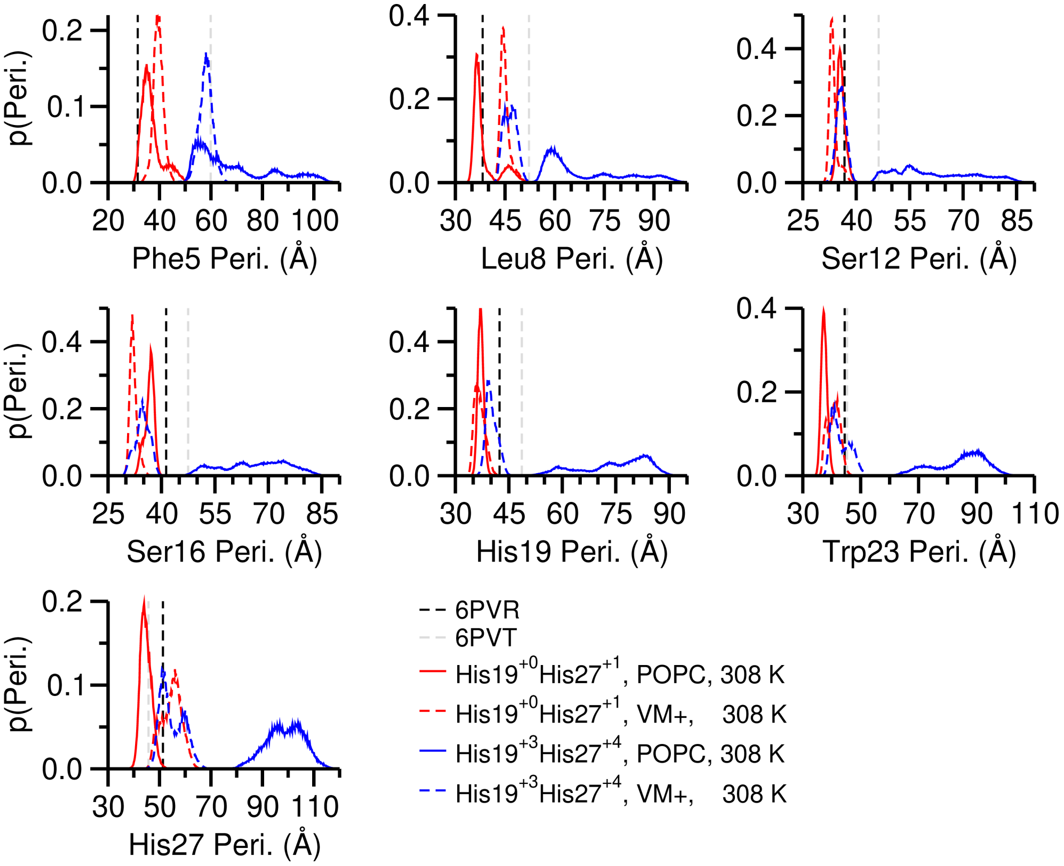


**Figure S12. Impact of fluid-phase lipid compositions on perimeters of the pore-lining residues.** Probability distributions of perimeters of the pore-lining residues in the closed (red lines) and open (blue lines) FPS simulations with POPC (solid lines) and VM+ (dashed lines) bilayers at 308.15 K. A perimeter was computed for the quadrilateral connecting the $C_\alpha$ atoms of four symmetric residues. The vertical black and grey dashed lines represent the closed (PDB ID: 6PVR) and open (PDB ID: 6PVT) ssNMR structures, respectively.

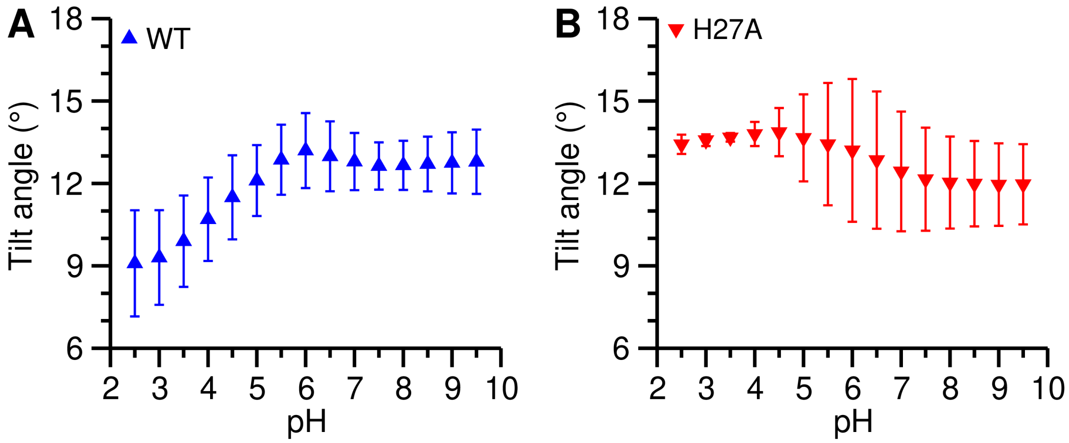


**Figure S13. Tilt angle between the N-terminal helical bundle (residues 7–14) and the Z-axis in the WT (A) and H27A mutant (B) CpHMD simulations.** Calculations were performed on the heavy atoms and insensitive to atom selections (Figure S14). Means over the simulation time were plotted versus pH values and errors were standard deviations estimated from block analysis.

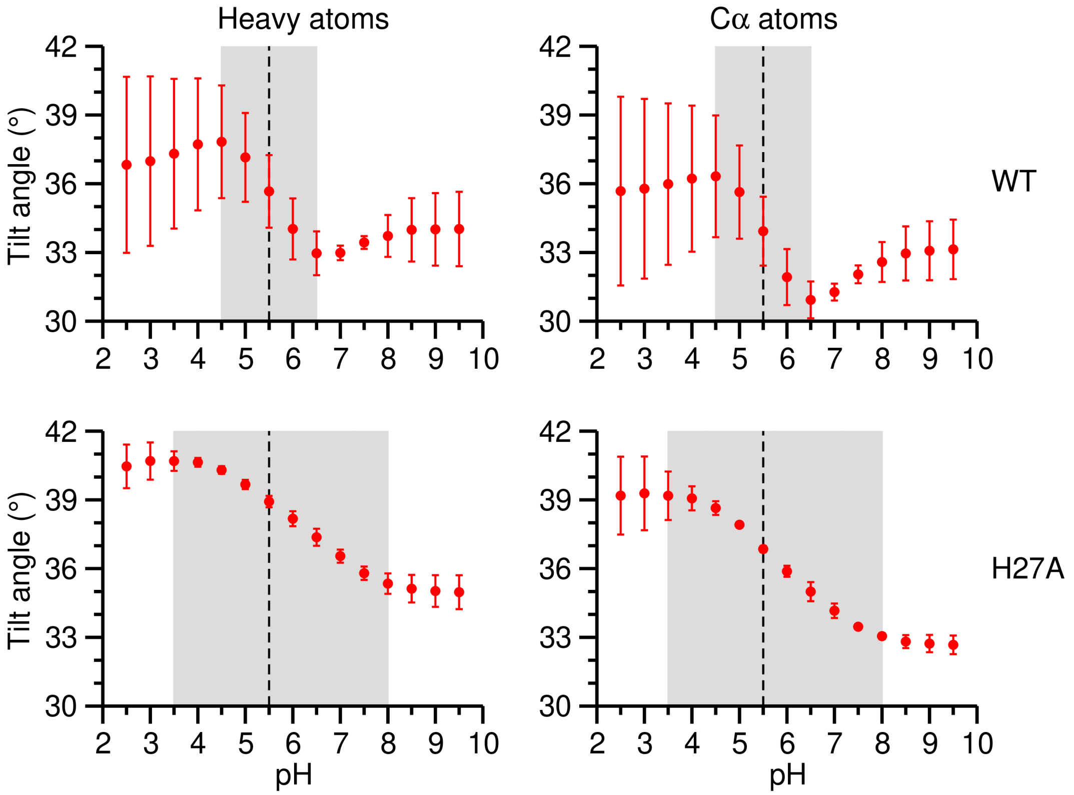


**Figure S14. Average tilt angle between the C-terminal helices (residues 15–28) and the N-terminal helical bundle (residues 7–14) in the WT (top) and H27A mutant (bottom) CpHMD simulations.** Means over the simulation time were plotted versus pH values and errors were standard deviations estimated from block analysis. Gray boxes and vertical black lines mark the transition regions and midpoints, respectively. Calculations using heavy or $C_\alpha$ atoms show consistent pH dependence, though heavy atoms yield 1–2° larger angles with smaller deviations.

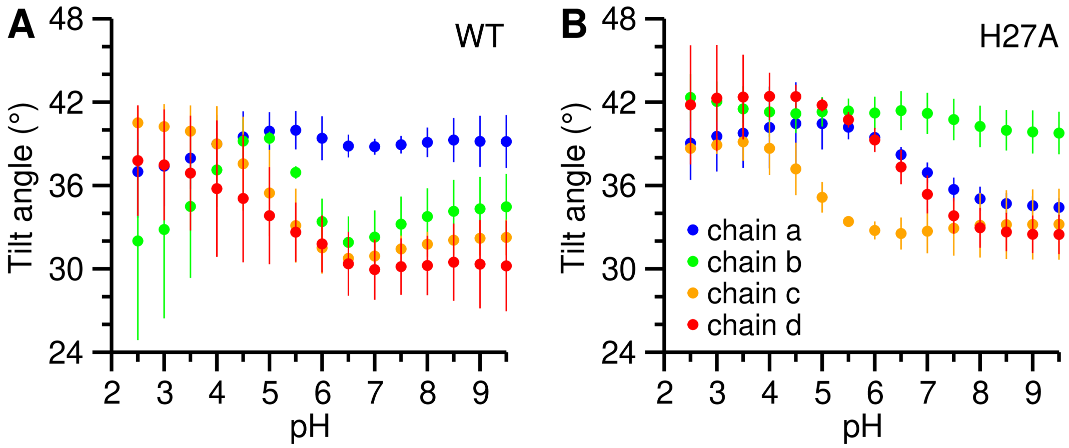


**Figure S15. Tilt angle between each C-terminal helix (residues 15–28) and the N-terminal helical bundle (residues 7–14) in the WT (A) and H27A mutant (B) CpHMD simulations.** Calculations were performed on the heavy atoms and insensitive to atom selections (Figure S14). Means over the simulation time were plotted versus pH values and errors were standard deviations estimated from block analysis.

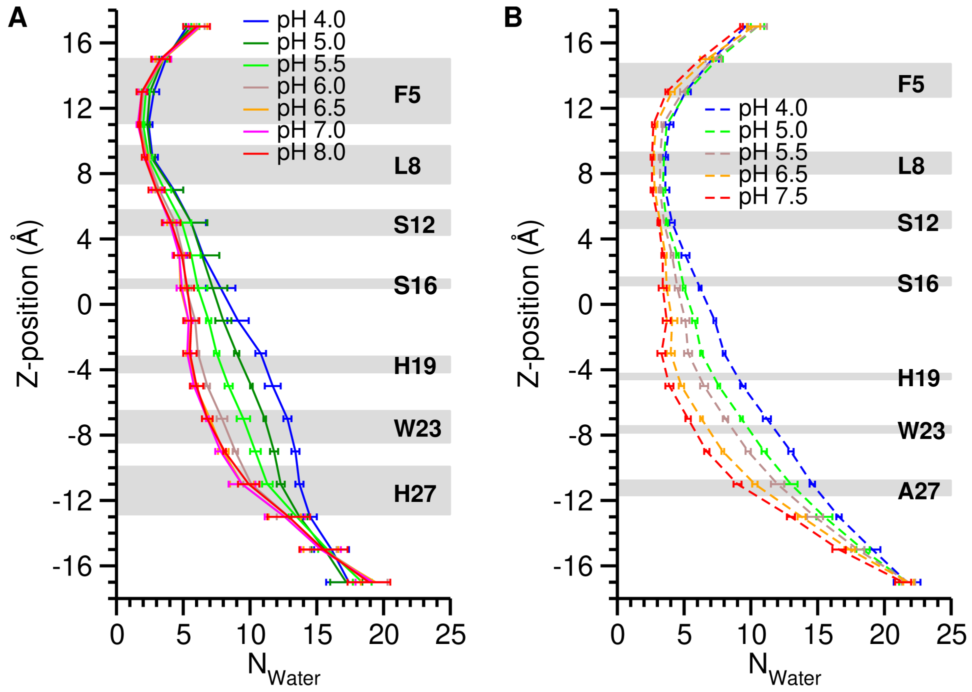


**Figure S16. pH-dependent average number of water molecules along the channel axis ($N_{Water}$) in the WT (A) and H27A mutant (B) simulations.** Water molecules were counted in 2-Å bins. Means over the simulation time were plotted versus pH values and errors were standard deviations estimated from block analysis. Gray boxes indicate the side-chain center-of-mass (COM) of the pore-lining residues (centroid and halved box width respectively represent the mean and standard deviation calculated from the plotted pH values). Note that the COMs were computed using heavy atoms and projected to the channel axis that centers the four Ala17 $C_\alpha$ atoms at the origin.

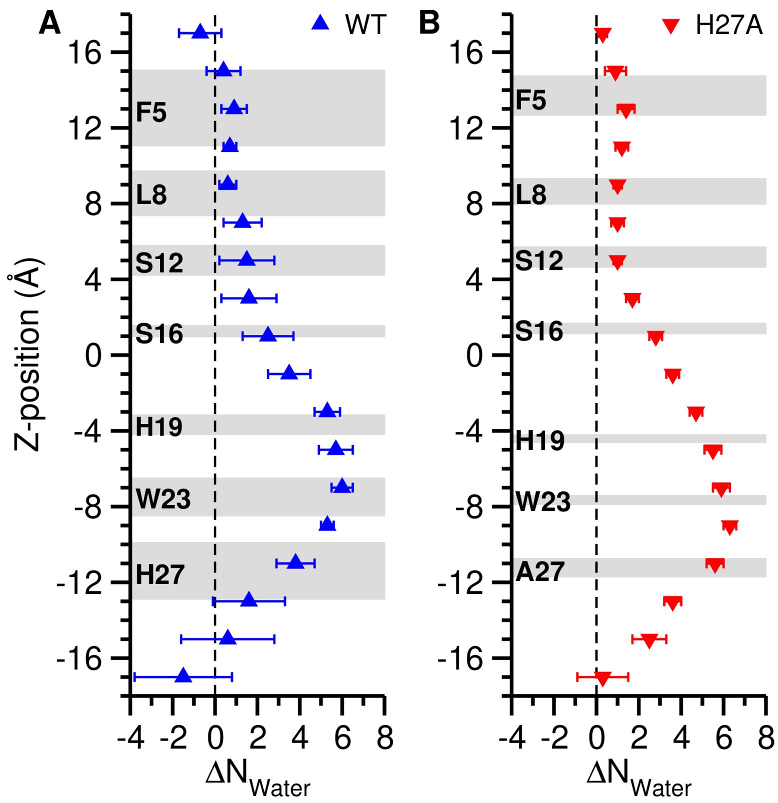


**Figure S17. Changes in the channel hydration upon activation (ΔN$_{Water}$) in the WT (A) and H27A mutant (B) CpHMD simulations.** ΔN$_{Water}$ was defined as the shift of the open-state (pH 4.0) N$_{Water}$ relative to the closed-state (pH 8.0 and 7.5 for WT and H27A, respectively) value. Errors were calculated through error propagation. Gray boxes indicate the side-chain center-of-mass (COM) of the pore-lining residues (centroid and halved box width respectively represent the mean and standard deviation). Vertical dashed lines mark ΔN$_{Water}$ of 0 $H_2O$ molecule.

**Table S6. Calculated microscopic/stepwise p$K_a$s of the GLU/HIS residues in BM2[a]**

| Residue[a] | | Simulation time window (ns) | | | | | | | |
|---|---|---|---|---|---|---|---|---|---|
| | | WT | | | | H27A | | | |
| | | 40–60 | 60–80 | 80–100 | 40–100 | 40–60 | 60–80 | 80–100 | 40–100 |
| E3 | $pK_a^a$ | 3.4 (0.9) | 3.5 (0.8) | 3.7 (0.8) | 3.5 (0.8) | 3.3 (0.8) | 3.2 (0.9) | 3.4 (0.8) | 3.3 (0.8) |
| | $pK_a^b$ | 3.5 (0.9) | 3.8 (0.8) | 3.7 (0.8) | 3.6 (0.8) | 3.5 (0.9) | 3.3 (0.8) | 3.5 (0.8) | 3.4 (0.8) |
| | $pK_a^c$ | 3.7 (0.8) | 3.7 (0.8) | 3.5 (0.8) | 3.6 (0.8) | 3.4 (0.9) | 3.7 (0.8) | 3.6 (0.8) | 3.5 (0.8) |
| | $pK_a^d$ | 3.5 (0.8) | 3.5 (0.8) | 3.5 (0.8) | 3.5 (0.8) | 3.5 (0.7) | 3.6 (0.8) | 3.5 (0.9) | 3.5 (0.8) |
| H19 | $pK_1$ | 6.9 (0.8) | 6.8 (0.7) | 7.0 (0.7) | 6.9 (0.7) | 7.1 (0.9) | 7.0 (0.9) | 7.0 (0.9) | 7.0 (0.9) |
| | $pK_2$ | 5.9 (0.8) | 6.1 (0.7) | 6.0 (0.7) | 6.0 (0.7) | 6.1 (0.9) | 6.2 (0.9) | 6.2 (0.9) | 6.1 (0.9) |
| | $pK_3$ | 5.5 (0.8) | 5.3 (0.8) | 5.4 (0.8) | 5.4 (0.8) | 5.2 (0.9) | 5.3 (0.9) | 5.4 (0.9) | 5.3 (0.9) |
| | $pK_4$ | 4.8 (0.9) | 4.7 (0.8) | 4.7 (0.7) | 4.7 (0.8) | 4.7 (0.9) | 4.6 (0.9) | 4.5 (0.9) | 4.6 (0.9) |
| H27 | $pK_1$ | 7.4 (0.8) | 7.5 (0.7) | 7.6 (0.8) | 7.5 (0.8) | - | - | - | - |
| | $pK_2$ | 6.9 (0.8) | 6.9 (0.8) | 6.9 (0.9) | 6.9 (0.8) | - | - | - | - |
| | $pK_3$ | 6.2 (0.8) | 6.3 (0.7) | 6.1 (0.8) | 6.2 (0.7) | - | - | - | - |
| | $pK_4$ | 5.3 (0.8) | 5.3 (0.6) | 5.3 (0.7) | 5.3 (0.6) | - | - | - | - |

[a] p$K_a$s were calculated within specific simulation windows. Parenthesized are the Hill coefficients *n*.

##### 4.2 Figures and tables discussed in the supporting information

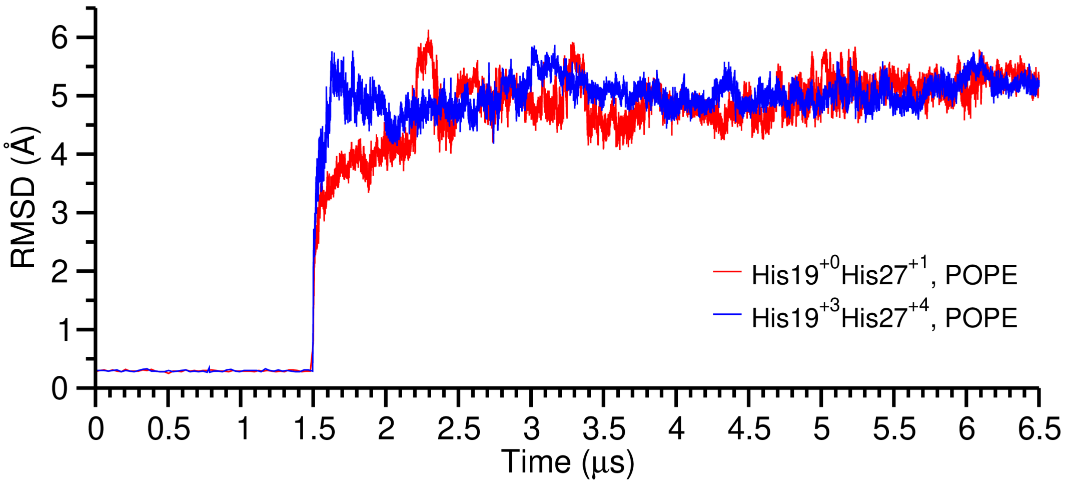


**Figure S18. Backbone RMSDs of the WT BM2 in gel-phase bilayers.** Time series of RMSD in the closed (red line) and open (blue line) FPS simulations with POPE bilayers at 280.15 K. Sampled every 100 ps.
